# CryoDRGN: Reconstruction of heterogeneous structures from cryo-electron micrographs using neural networks

**DOI:** 10.1101/2020.03.27.003871

**Authors:** Ellen D. Zhong, Tristan Bepler, Bonnie Berger, Joseph H. Davis

## Abstract

Cryo-EM single-particle analysis has proven powerful in determining the structures of rigid macromolecules. However, many protein complexes are flexible and can change conformation and composition as a result of functionally-associated dynamics. Such dynamics are poorly captured by current analysis methods. Here, we present cryoDRGN, an algorithm that for the first time leverages the representation power of deep neural networks to efficiently reconstruct highly heterogeneous complexes and continuous trajectories of protein motion. We apply this tool to two synthetic and three publicly available cryo-EM datasets, and we show that cryoDRGN provides an interpretable representation of structural heterogeneity that can be used to identify discrete states as well as continuous conformational changes. This ability enables cryoDRGN to discover previously overlooked structural states and to visualize molecules in motion.

## Main

Single particle cryo-electron microscopy (cryo-EM) is a rapidly maturing method for high-resolution structure determination of large macromolecular complexes^1,2^. Major advances in hardware^3-5^ and software^4-9^ have streamlined the collection and analysis of cryo-EM datasets, such that structures of rigid macromolecules can routinely be solved at near atomic resolution^10,11^. However, a major computational bottleneck remains when conformational or compositional heterogeneity is present in the sample.

The crux of cryo-EM structure determination is the computational task of reconstruction, where algorithms must learn the 3D density or densities from the recorded dataset of 2D particle images^12^. While the standard formulation of reconstruction assumes that each 2D image is generated from a single, static structure, in reality, each image contains a unique snapshot of the molecule of interest. While this heterogeneity complicates reconstruction, it presents an opportunity for single particle cryo-EM to reveal the conformational landscape of dynamic macromolecular complexes.

Existing tools for heterogeneous reconstruction often make strong assumptions on the type of heterogeneity in the dataset. Most commonly, heterogeneity is modeled as though it originates from a small number of independent, *discrete* states^13-16^, consistent with molecules undergoing cooperative conformational changes. However, because the number of underlying structural states are unknown, such discrete classification approaches are error-prone and often result in the omission of potentially relevant structures. Moreover, this approach fails to model molecules that undergo continuous conformational changes. In these conformationally heterogeneous systems, user-defined masks have been employed to resolve isolated rigid-body motions^17^, but these approaches require assumptions on the types and location of molecular motions. Additionally, new theoretical methods have been proposed to model global continuous heterogeneity^18-20^, however no such tools have been made available.

Here, we present cryoDRGN (Deep Reconstructing Generative Networks), a cryo-EM reconstruction method that uses a deep neural network to directly approximate the molecule’s continuous 3D density function (**Fig. 1a**). We designed this tool based on the reasoning that deep neural networks, which are known for their ability to model continuous, nonlinear functions, might effectively capture dynamical structures. We show that this neural network representation of structure, which we call a *deep coordinate network*, can efficiently learn heterogeneous ensembles of high-resolution structures from single particle cryo-EM datasets^21^.

**Figure 1.**
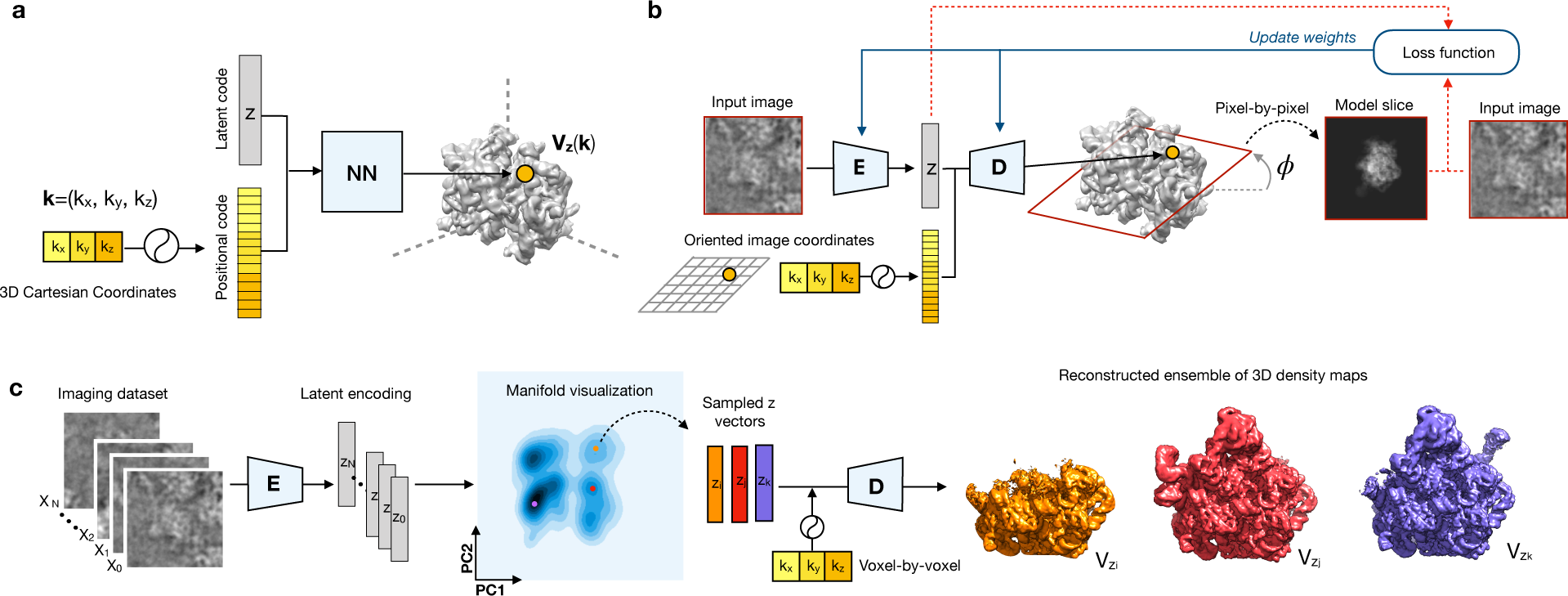
The cryoDRGN method for heterogeneous single particle cryo-EM reconstruction. **a)** A *deep coordinate network* approximates a molecule’s density as a function of featurized 3D Cartesian coordinates and continuous latent variables, *z*, which define a continuous manifold of heterogeneous structures. **b)** The overall cryoDRGN training framework consists of two neural networks structured in an encoder/decoder architecture. Data is represented in the Fourier domain in order to relate 2D images as slices out of the 3D density map. During training, an input image is encoded in latent space by the encoder network (E). A 2D lattice is rotated by the image’s previously determined pose, *ϕ*, to represent the 3D coordinates of the image slice. Given the coordinates and a sample of the predicted latent variable *z*, the image is reconstructed pixel-by-pixel through the decoder (D), *i*.*e*. the deep coordinate network. The loss function is a variational upper bound on the data likelihood and consists of the image reconstruction error and latent loss (red arrows), which is used to update neural network weights by stochastic gradient descent (blue arrows). **c)** After training, the encoder can be used to visualize the dataset’s distribution in latent space (manifold visualization), and the decoder can be used to directly reconstruct structures at arbitrary points from the latent code. Example micrographs and reconstructed density maps from EMPIAR 10076^32^.

To learn this representation, cryoDRGN introduces an image-encoder/volume-decoder framework to learn a latent representation of heterogeneity from single particle cryo-electron micrographs. Once trained, users can visualize the dataset in the low-dimensional latent space, which we find reflects the structural heterogeneity of the imaged molecule. This structural heterogeneity can then be interrogated by generating 3D density maps at an arbitrary number of desired positions within the latent space, which can be used to visualize continuous structural trajectories.

CryoDRGN is a powerful and general approach for analyzing heterogeneity in imaging datasets and can be used to reconstruct both compositionally and conformationally heterogeneous structures. We demonstrate its efficacy by reconstructing and analyzing structures of the eukaryotic ribosome, the assembling bacterial ribosome, and the pre-catalytic spliceosome complex. In these machines, we discover new conformational states and observe dynamic molecular motions. CryoDRGN is distributed as an open-source tool^22^ that can be easily integrated in existing pipelines and is freely available at cryodrgn.csail.mit.edu.

## Results

### CryoDRGN architecture and training

CryoDRGN performs heterogeneous reconstruction by learning a neural network representation of 3D structure from single particle cryo-EM micrographs. In contrast to traditional reconstruction algorithms, which represent the 3D density map on a discretized voxel array, cryoDRGN uses a neural network to predict density as a function of 3D Cartesian coordinates. We call this architecture^23-25^ a *deep coordinate network*. To model heterogeneity, the deep coordinate network can be extended to predict density as a function of both 3D coordinates and continuous latent variables, *z*, which define a n-dimensional manifold of heterogeneous structures (**Fig. 1a**). Coordinates are featurized with a positional encoding function before they are input to the deep coordinate network (Methods). This choice of model assumes that structures can be embedded within a continuous low-dimensional space, *i*.*e*. the latent space, where the dimensionality of the latent space is defined by the user.

To train this neural network representation of 3D structure from a single particle cryo-EM dataset, we develop an encoder–decoder architecture based on the Variational Autoencoder (VAE)^26,27^ (**Fig. 1b**). The structure is represented in the Fourier domain in order to relate 2D images as central slices out of a density map according to the Fourier slice theorem^28^. For a given image, *X*, the encoder neural network predicts a distribution of possible latent variable values, *q*(*z*|*X*). The image’s oriented 3D pixel coordinates are computed from the image’s pose assignment provided by a previously determined consensus reconstruction. Then, given a sample from the encoder distribution *z* ∼ *q*(*z*|*X*) and the 3D coordinates of the slice, the image is reconstructed pixel-by-pixel through the deep coordinate network. The networks are trained jointly using an objective function that seeks to optimize a variational upper bound on the data likelihood as in standard VAEs^25^. This objective function consists of the image reconstruction error and a regularization term on the latent space. The parameters of the neural networks are iteratively updated by gradient descent on this objective function.

After training, the encoder network is used to map images into the low-dimensional latent space, where we define 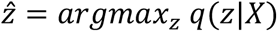 as each image’s “latent encoding” (**Fig. 1c**). The full distribution of latent encodings can then be visualized to study the latent space data manifold. To explore the ensemble of structures, the deep coordinate network can directly reconstruct a 3D density map given a desired value of the latent variable *z* and the 3D coordinates of a voxel array.

### Deep coordinate networks can learn static structures from homogeneous datasets

To test the efficacy of neural networks in representing 3D structure, we first trained a deep coordinate network with no latent variable input to learn the homogenous structure of the *Plasmodium falciparum* 80S (*Pf*80S) ribosome from the EMPIAR 10028 dataset^29^. The network was trained on full resolution images (D=360, Nyquist limit of 2.7 Å), where image poses were obtained from a consensus reconstruction in cryoSPARC^30^. We found that the deep coordinate network produced a structure qualitatively matching the traditional reconstruction (**Fig. 2a**) at resolutions up to ∼4.0 Å at an FSC=0.5 threshold (**Fig. 2b**).

**Figure 2.**
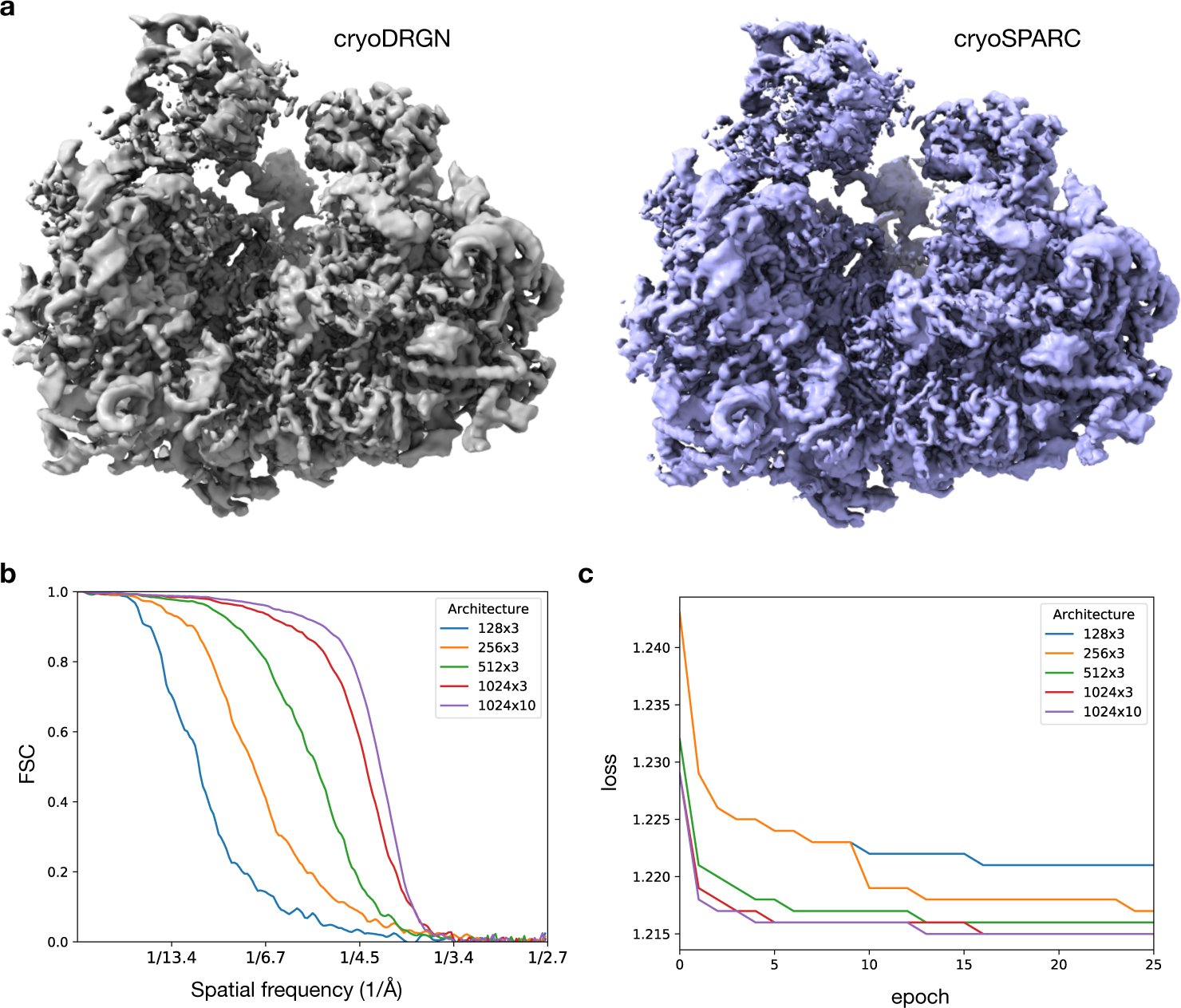
Deep coordinate network representation of static structure. **a)** Reconstructed density map produced by a deep coordinate network with 10 hidden layers of dimension 1024 trained on particle images from EMPIAR 10028^29^ (D=360, Nyquist limit of 2.7 Å) and a traditional homogeneous reconstruction in cryoSPARC^29^. **b)** Fourier shell correlation (FSC) curves between the density map produced by deep coordinate networks of varying architectures (nodes x hidden layers) and the traditional homogeneous reconstruction in (a) after 25 epochs of training. **c)** Average loss over the dataset during training deep coordinate networks of varying architectures on EMPIAR 10028^29^.

As neural networks have a fixed capacity for representation that is constrained by their architecture, we compared architectures of different sizes to evaluate the tradeoff between representation power and training speed. We found that larger architectures converged to lower values of the objective function (**Fig. 2c**) and correlated with the traditionally reconstructed map at higher resolution (**Fig. 2b**). These improvements in the resulting structure came at the cost of extended training times, suggesting that the architecture and the image size should be tuned to suit the desired balance of training speed and achievable resolution (**Supplementary Fig. 1**).

### CryoDRGN models both discrete and continuous structural heterogeneity

We next used simulated single particle cryo-EM datasets to test if the complete cryoDRGN framework could perform *heterogeneous* reconstruction. This simulation-based approach allowed us to quantitatively evaluate the method’s performance by comparing the reconstructed density maps to the ground truth density maps. To simulate continuous motions, we constructed an atomic model of a hypothetical protein complex and iteratively rotated one bond’s dihedral angle, resulting in a series of 50 distinct but closely-related atomic models. We then generated density maps along this reaction coordinate to serve as the ground truth density maps (**Fig. 3a**). Cryo-EM micrographs were generated by projecting the ground truth maps with random poses, followed by application of the contrast transfer function (CTF) and the addition of noise (see Methods). To simulate a compositionally heterogeneous dataset, this procedure was repeated by mixing images generated from the bacterial 30S, 50S, and 70S ribosomal density maps (**Fig. 3d**). The cryoDRGN networks were then provided these simulated images and their corresponding poses, and were trained with 1-dimensional (1D) latent variable models.

**Figure 3.**
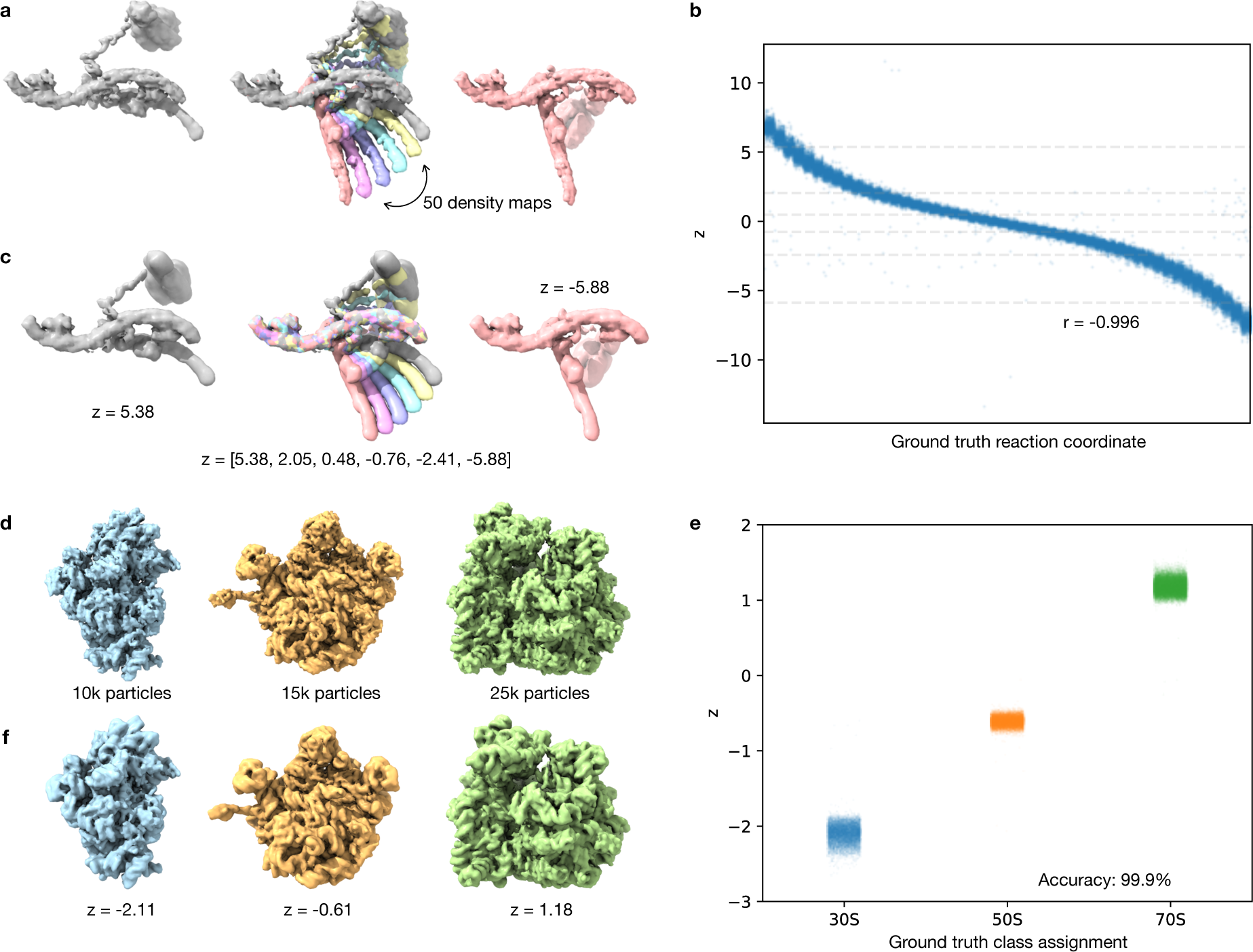
CryoDRGN heterogeneous reconstruction of simulated datasets with continuous and discrete heterogeneity. **a)** Ground truth density maps sampled along a reaction coordinate that describes the transition from the leftmost to rightmost structure used to simulate a dataset with continuous heterogeneity. **b)** Predicted latent encoding for each image of the dataset from (a) after training a cryoDRGN 1D latent variable model versus the ground truth reaction coordinate describing the motion (Spearman *r* = −0.996). **c)** Reconstructed structures at specified values of the latent variable, shown as dotted lines in (b). **d)** Ground truth density maps of the bacterial 30S, 50S, and 70S ribosome used to simulate a dataset with discrete heterogeneity. **e)** Predicted latent encoding for each particle image of (d) variable after training a cryoDRGN 1D latent variable model vs. its ground truth class assignment (classification accuracy of 99.9%). **f)** Reconstructed structures at specified values of the latent variable from (e).

When trained on the dataset with continuous heterogeneity, we found that cryoDRGN accurately modeled the full continuum of structures as assessed by two criteria. First, the latent encoding of each image produced by the encoder network correlated well with the dihedral angle of the underlying model (Spearman *r* = −0.996), which we characterize as the ground truth reaction coordinate (**Fig. 3b**). Second, when provided a series of latent variable values, the deep coordinate network produced structures that correlated with the ground-truth maps (**Fig. 3c**). We note that the deep coordinate network can generate an arbitrary number of conformations along the trajectory, and found that for 100 images equally spaced along the reaction coordinate, the generated structure at each image’s predicted latent encoding correlated well with its ground truth map (**Supplementary Fig. 2**).

When cryoDRGN was trained on the compositionally heterogeneous dataset, we observed that the encoder network mapped particles to three distinct clusters in latent space (**Fig. 3e**). These clusters aligned with the ground truth class assignments from the 30S, 50S, and 70S ribosome (classification accuracy of 99.9%), and the appropriate ribosomal structures were generated by the deep coordinate network when provided with latent variable values at the corresponding cluster centers (**Fig. 3f, Supplementary Fig. 2**).

### CryoDRGN uncovers residual heterogeneity in a high-resolution cryo-EM reconstruction

We next evaluated cryoDRGN’s ability to learn heterogeneous structures from real cryo-EM data, which contains structured noise and imaging artifacts that are difficult to simulate. When analyzing a homogeneous reconstruction of the *Pf*80S ribosome, Wong *et al*. observed flexibility in the small subunit head region and missing density for peripheral rRNA expansion segment elements that prevented completion of an atomic model in these regions^29^. To explore if this unresolved density resulted from residual heterogeneity, we trained a 10-dimensional (10D) latent variable model with cryoDRGN on their deposited dataset (EMPIAR-10028), using poses from a consensus reconstruction in cryoSPARC^30^. We then visualized the dataset’s latent encodings using principal component analysis (PCA) (**Fig. 4a**) and observed a subset of particles separated along PC1. A density map generated by the deep coordinate network from this region of latent space revealed a distinct conformation of the 40S subunit, which was rotated relative to the 60S subunit (**Fig. 4b,c**). Concomitant with the inter-subunit rotation, we observed the disappearance of the inter-subunit bridge formed by the C-terminal helix of eL8, which is consistent with Sun *et al*.’s characterization of Pf80S dynamics^31^. We further explored structural heterogeneity by performing *k*-means clustering of the latent encodings with *k*=20 clusters and subsequently generating structures at the cluster centers. We observed diverse structures including those bearing a rotated *Pf*40S head group, those missing the head group, and those with clearly resolved rRNA helices that were absent from the homogeneous reconstruction (**Fig. 4c**).

**Figure 4.**
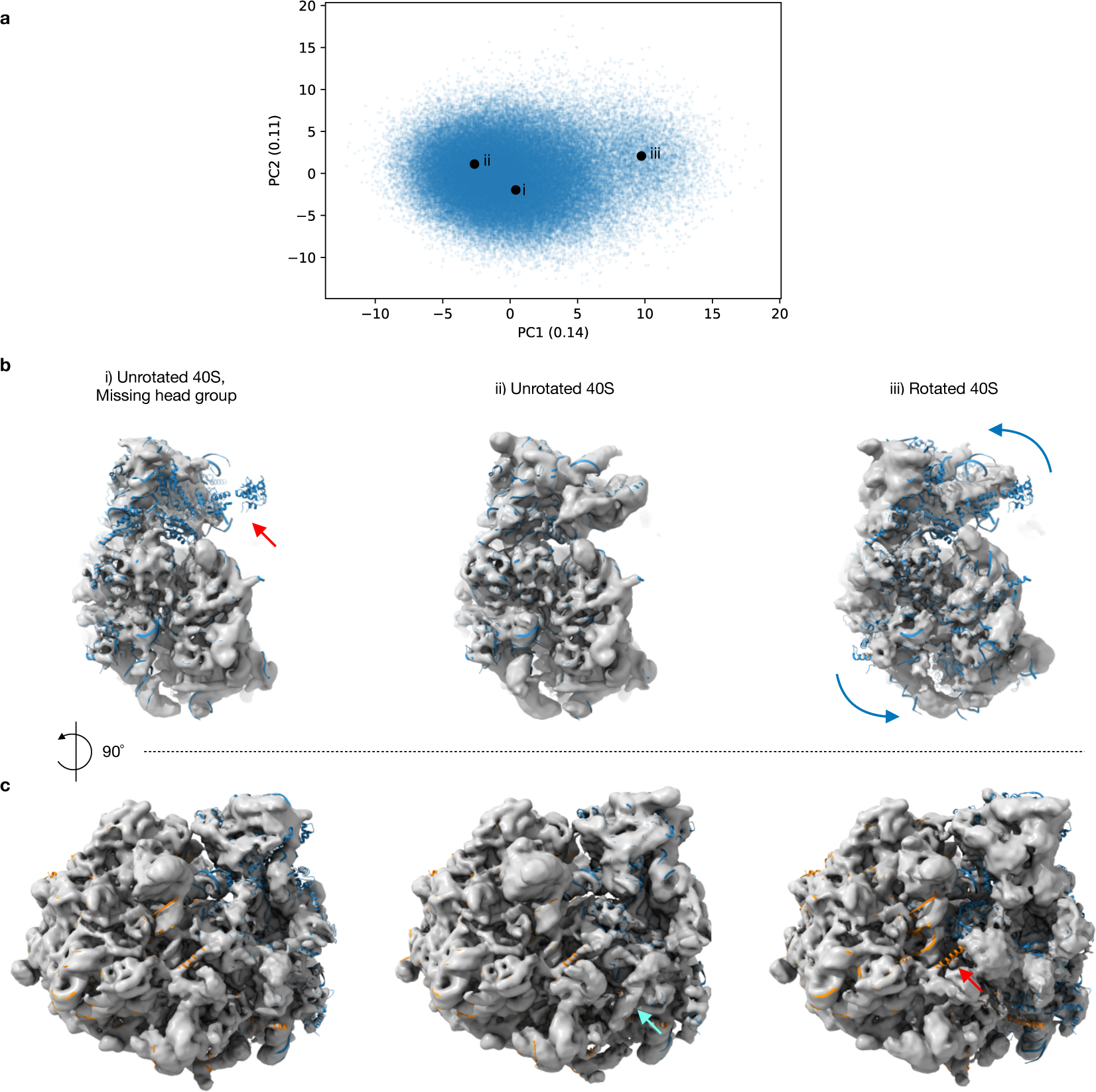
CryoDRGN heterogeneous reconstruction of the *Pf*80S ribosome. **a)** PCA projection of latent space encodings after training a 10D latent variable model on particle images from EMPAIR-10028^29^. **b)** Three representative density maps that were reconstructed at the points depicted in (a) are shown with a docked atomic model (PDB 3J79, 3J7A) of the 40S (blue). The red arrow highlights the missing 40S head group, and the blue arrow depicts the rotation of the 40S relative to the 60S. **c)** Additional views of the structures shown in (b), with atomic model of the 60S colored in orange. The cyan arrow notes the presence of an additional RNA helix not present in the homogeneous reconstruction, and the red arrow notes the disappearance of the C-terminus of eL8 in the rotated state.

### CryoDRGN automatically partitions assembly states of the bacterial ribosome

Next, we assessed cryoDRGN’s ability to analyze and reconstruct density maps from a dataset known to contain substantial compositional and conformational heterogeneity. For this assessment, we investigated a highly heterogeneous mixture of assembly intermediates of the *E. coli* large ribosomal subunit (LSU). This dataset (EMPIAR 10076) had previously been analyzed through multiple expert-guided rounds of hierarchical 3D classification resulting in 13 discrete structures that were grouped into 4 major classes^32^. These particles were obtained by crudely fractionating a lysate with the explicit goal of imaging and later analyzing the full ensemble of cellular assembly intermediates. As such, a substantial fraction of the published particle stack corresponds to non-ribosomal impurities that were discarded during 3D classification in the original analysis (26,575 out of 131,899 images). Despite this heterogeneity, a homogeneous reconstruction of the full dataset produced a consensus structure of the mature LSU (GSFSC resolution of 3.2 Å), suggesting that even in the presence of these impurities, the heterogeneous ribosomal particles could be aligned to the rigid core of the LSU, which enabled analysis using cryoDRGN.

To assess the degree of heterogeneity in the data, we first trained a 1D latent variable model on down-sampled images (D=128, Nyquist limit of 6.6 Å) using image poses from a consensus reconstruction. After model training, the encoder network was used to predict the latent encoding for each particle, and the resulting histogram of the full dataset’s encodings revealed five distinct peaks. Four of the peaks corresponded to each of the four major classes of the LSU, and the fifth peak near *z* = −2 captured particles that were unassigned by Davis *et al*. (**Fig. 5a**)^32^. This clear separation in latent space suggested that cryoDRGN can identify sample impurities without supervision. When using the subset of particles from this region (assigned *z* ≤ −1), neither 2D class averages nor a traditional 3D reconstruction produced structures consistent with assembling ribosomes (**Supplementary Fig. 3**). As we do not wish to model these impurities, we filtered the dataset by the latent variable, keeping 101,604 images with *z* > −1 for further analysis.

**Figure 5.**
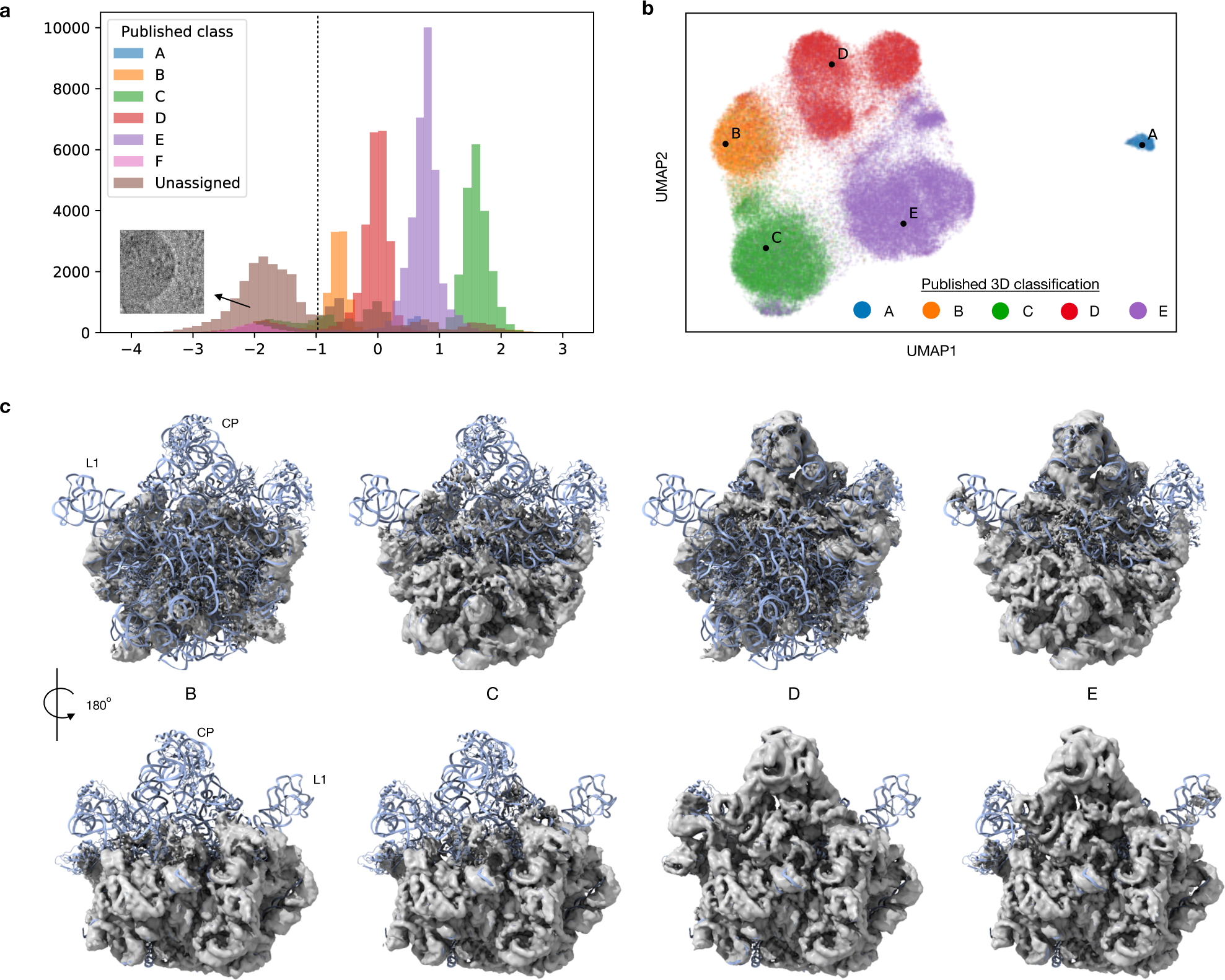
CryoDRGN heterogeneous reconstruction of the assembling large ribosomal subunit from *E. coli*. **a)** Histograms of latent encodings of particle images from EMPIAR 10076^32^ after training a cryoDRGN 1D latent variable model. Overlaid histograms are shown for particles from each published major class assignments from *Davis et al*^31^. A cutoff of *z* = −1 was used to filter impurities from the dataset for subsequent analyses. Example image of an ice artifact predicted at *z* = −2. **b)** UMAP visualization of latent encodings after training a cryoDRGN 10D latent variable model, colored by the published major class assignments^32^. **c)** CryoDRGN reconstructed density maps of the major assembly states of the LSU generated from points B-E shown in (b) along with a docked atomic model (PDB 4YBB).

To explore the heterogeneity within these major assembly states, we trained a 10D latent variable model on the remaining high-resolution images (D=256, Nyquist limit of 3.3 Å). We visualized the resulting 10D latent encodings using UMAP^33^, and observed particle super-clusters corresponding to major classes of LSU assembly (**Fig. 5b**) and sub-clusters that aligned with Davis *et al*.’s minor class assignments (**Supplementary Fig. 4**)^32^. We found that when provided latent codes from these clusters, the decoder network generated structures matching the major (**Fig. 5c**) and minor (**Supplementary Fig. 5**) assembly states of the LSU. With the 10D latent variable model, we also noted a clearly separated cluster of particles assigned to class A, and structures sampled from this region of latent space reconstructed the 70S ribosome, an impurity in the dataset (**Supplementary Fig. 6**). Finally, we identified a small cluster of ∼1,200 particles in latent space adjacent to the class C cluster whose particles were classified into class E by Davis *et al*. (**Supplementary Fig. 4**). The density map reconstructed by the deep coordinate network from this region revealed a previously unreported assembly intermediate that we newly call class C4. Like the other class C structures, class C4 lacked the central protuberance, but bore clearly resolved density for helix 68, which was only present in classes E4 and E5 from Davis *et al*^32^. Traditional voxel-based back-projection of the particle images constituting this cluster reproduced a similar, albeit lower-resolution structure, confirming the existence of this structural state in the original dataset (**Supplementary Fig. 6**).

### CryoDRGN reveals dynamic continuous motions in the pre-catalytic spliceosome

Finally, we evaluated the performance of cryoDRGN in analyzing micrographs of the pre-catalytic spliceosome (EMPIAR 10180)^34^. Plaschka *et al*. employed extensive expert-guided focused classifications to reconstruct a composite map for this complex and suggested that the complex sampled a continuum of conformations^34^. To understand how cryoDRGN would encode such continuous structural heterogeneity in latent space, we first trained a 10D latent variable model on the downsampled images (D=128, Nyquist limit of 8.5 Å) using image poses derived from a consensus reconstruction. Multiple clusters were observed in the latent space encodings (**Fig. 6a**). After sampling structures from the latent space, we observed expected spliceosome conformations from the largest cluster, poorly resolved structures from the leftmost cluster, structures lacking density for the SF3b domain from a third cluster, and additional density consistent with particle aggregation from the uppermost cluster (**Fig. 6b**). To focus our analysis on bone-fide pre-catalytic spliceosome particles, we leveraged the latent space representation to eliminate any particles that mapped to the undesired clusters from two replicate runs, which resulted in a final particle stack of 150,098 images.

**Figure 6.**
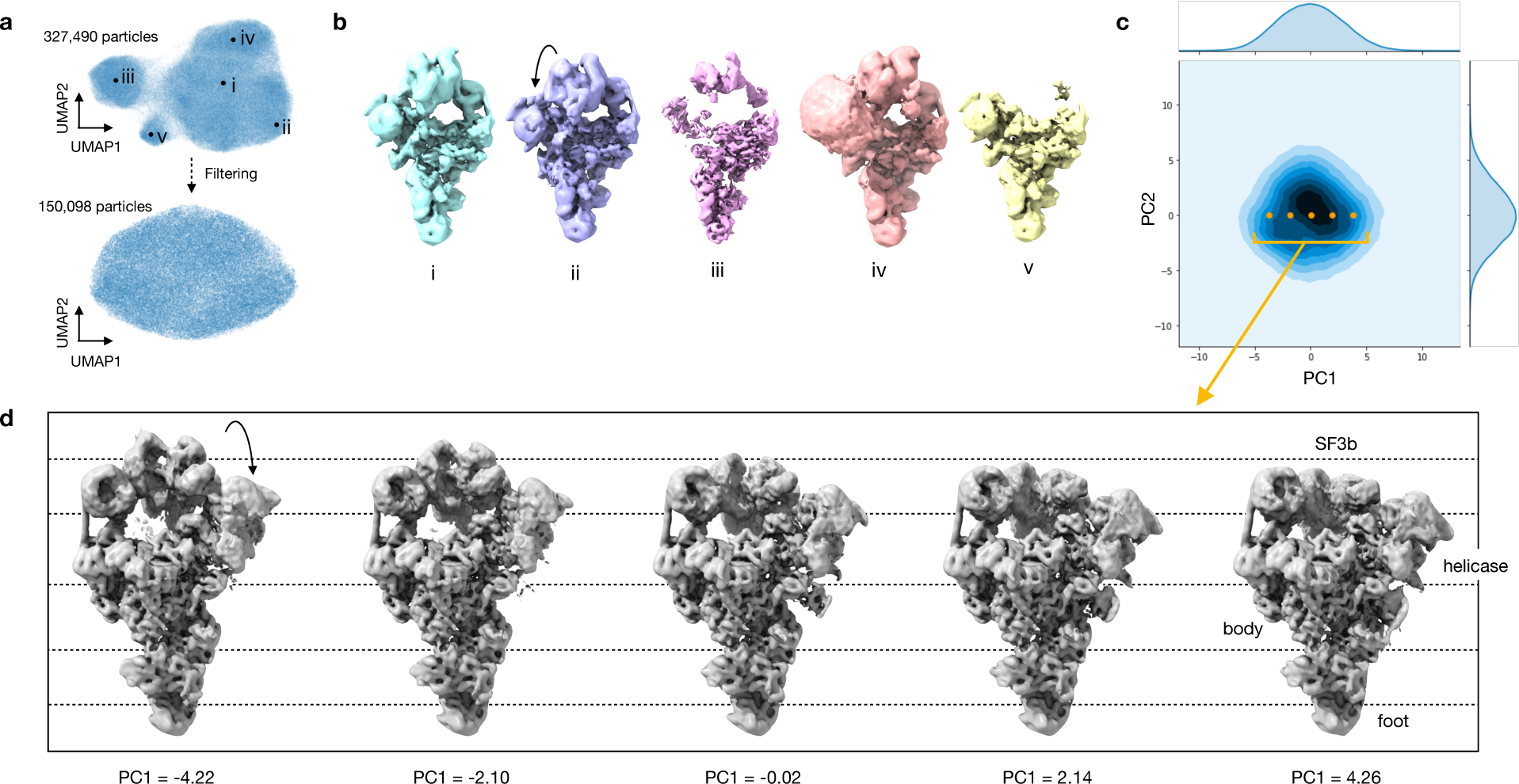
CryoDRGN heterogeneous reconstruction of the pre-catalytic spliceosome. **a)** UMAP visualization of latent encodings after training a 10D latent variable model with cryoDRGN on EMPIAR 10180^33^, before (top) and after (bottom) particle filtering. **b)** Representative structures generated at points shown in (a) which depict the expected structures (i,ii), broken particles (iii), particles with apparent aggregation (iv), and the complex lacking the SF3b domain (v). **c)** PCA projection of latent space encodings after training a 10D latent variable model on the filtered images. **d)** Structures generated by traversing along PC1 of the latent space encodings at points shown in (c).

With the filtered particle stack, we trained a 10D model on higher resolution images (D=256, Nyquist limit of 3.4 Å), and visualized the dataset’s latent encodings in 2D using PCA and UMAP (**Fig. 6a,c**). The visualized data manifold was unfeatured, consistent with a molecule undergoing non-cooperative conformational changes. By generating structures along the first principal component of the latent space encodings, we reconstructed a trajectory of the SF3b and helicase domains in motion (**Fig. 6d**), which smoothly transitioned from an elongated state to one compressed against the body of the spliceosome. A similar traversal along the second PC produced a continuous trajectory of the SF3b and helicase domains moving in opposition (**Supplementary Fig. 7**). This anticorrelated motion of the SF3b and helicase domains in PC2, together with their correlated motion in PC1, suggested that the two domains move independently in the imaged ensemble. Finally, although trajectories along latent space PCs provide a summary of the extent of variability in the structure, cryoDRGN can also generate structures at arbitrary points from the latent space. By traversing along the *k-*nearest neighbor graph of the latent encodings and generating structures at the visited nodes, cryoDRGN generated a plausible trajectory of the conformations adopted by the pre-catalytic spliceosome (**Supplemental Movie 1**), highlighting the potential of single particle cryo-EM to uncover the conformational dynamics of molecular machines.

## Discussion

This work introduces cryoDRGN, a new method using neural networks to reconstruct 3D density maps from heterogeneous single particle cryo-EM datasets. The power of this approach lies in its ability to represent heterogeneous structures without simplifying assumptions on the type of heterogeneity. In principle, cryoDRGN is able to represent any distribution of structures that can be approximated by a deep neural network, a broad class of function approximators^35^. This flexibility contrasts with existing methods that impose strong assumptions on the types of structural heterogeneity present in the sample. For example, traditional 3D classification assumes a mixture of discrete structural classes, whereas multibody refinement assumes conformational changes are composed strictly of rigid-body motions. Although these approaches have proven useful, they are inherently unable to model true structural heterogeneity and thus often introduce bias into reconstructions. In contrast, we empirically show that deep coordinate networks can model both discrete compositional heterogeneity and continuous conformational changes without the aforementioned assumptions. For example, by using this less biased approach, we discovered heterogeneous states of the *Pf*80S ribosome that were originally averaged out of the homogeneous reconstruction. When analyzing the assembling *E. Coli* LSU dataset, cryoDRGN learned an ensemble of LSU assembly states without *a priori* specification of the number of states as is required for 3D classification. Finally, when analyzing the pre-catalytic spliceosome, we found that the continuous conformational changes cryoDRGN reconstructed lack the rigid-body boundary artifacts introduced from multibody refinement’s mask-based approach^17^.

### Interpretation of the latent space

A key feature of cryoDRGN is its ability to provide a low-dimensional representation of the dataset’s heterogeneity, which is given by each particle’s latent encoding. Subject to optimization, cryoDRGN organizes the latent space such that structurally related particles are in close proximity. Thus, visualization of the distribution of latent encodings can be informative in understanding the structural heterogeneity within the imaged ensemble. In both simulated and real datasets we find that continuous motions are embedded along a continuum in latent space (**Fig. 3b, 6c**) and that compositionally distinct states manifest as clusters (**Fig. 3e, 5b**). This observation suggests an interpretation of the latent encodings as an approximate conformational landscape, with regions of high-particle occupancy corresponding to low-energy states, and regions of lower-particle occupancy denoting higher energy states. We note however that structures reconstructed from unoccupied regions will not in general correspond to true physical intermediates, as cryoDRGN optimizes the likelihood of the observed data and these intermediates are not observed. Finally, in real datasets, there may exist images that do not originate from the standard single particle image formation model, for example, false positives encountered during particle picking^9^. We demonstrated the utility of the latent space encodings in identifying such impurities, ice artifacts, and other such out-of-distribution particles that may be filtered out in subsequent analyses (**Fig. 5a, 6a**).

### Visualizing structural trajectories

In addition to encoding particles in an unsupervised manner, cryoDRGN can reconstruct 3D density maps from user-defined positions in latent space. Because cryoDRGN learns a generative model for structure, an unlimited number of structures can be generated and analyzed, thus enabling visualization of structural trajectories. By leveraging the latent encodings of the particle images, users can directly traverse the data manifold and only sample structures from regions of latent space with significant particle occupancy. Indeed, we applied a well-established graph-traversal algorithm^36^ to visualize a data-supported path of the *Pf*80S ribosome, bL17-independent assembly of the bacterial ribosome, and the pre-catalytic spliceosome (**Supplemental Movies 1,2,3,4**).

### Practical considerations in choosing training hyperparameters

Although this method emphasizes an unsupervised approach to analyzing structural heterogeneity, cryoDRGN does require that the user define the dimensionality of the latent space and the architecture of both the encoder and decoder networks. We find that in practice, a 1D latent space is effective at distinguishing bona-fide particles from contaminants and imaging artifacts (**Fig. 5a**), and we recommend users initially employ such a model to filter their dataset. Additionally, we find that in our tested datasets, a 10D latent space provides sufficient representation capacity to effectively model structural heterogeneity, and that this 10D space can be readily visualized with PCA or UMAP. Notably, we recommend the use of such as 10D latent space instead of lower dimensional space as we have found that 10D spaces result in much more rapid overall training, which is consistent with similar observations of related overparameterized neural network architectures^37,38^. Finally, users must specify the number of nodes and layers in the neural networks. Here, we find an inverse relationship between neural network size and the achievable resolution of a given structure (**Supplemental Fig. 1**). Training larger networks on larger images is significantly slower, and we recommend that users perform an initial assessment using down-sampled images and relatively small networks before proceeding to high-resolution reconstructions.

### Discovering new states using cryoDRGN

CryoDRGN can be used to identify novel clusters of structurally-related particles, which can then be visualized by sampling a 3D structure from that region of latent space. Indeed, in analyzing the bL17-depleted LSU assembly dataset, we noted a completely new structural class, which like the C-classes, lacked the central protuberance, but like the most mature E classes, clearly bore a functionally critical inter-subunit helix (h68). This state was completely missed in traditional hierarchical classification^32^, and provides structural evidence that this vital intersubunit helix can dock in a native conformation in the absence of the central protuberance (**Supplementary Fig. 6**). Notably, we could validate the existence of this class by performing traditional back-projection using ∼1,000 particles from this cluster (**Supplementary Fig. 6**).

In future work, we envision using cryoDRGN to reveal the number of discrete classes, their constituent particles, and to produce initial 3D models that could be used as inputs for a traditional 3D reconstruction. Given the mature state of such tools^39,40^, this unbiased classification approach followed by traditional homogeneous reconstruction, particle polishing, and higher order image aberration correction, has the potential to produce very high-resolution structures of the full spectrum of discrete structural states without the need for expert-guided classification.

### Fully unsupervised 3D reconstruction

As implemented, cryoDRGN uses pose estimates resulting from a traditional consensus 3D reconstruction. In analyzing three publicly available datasets, we found that such consensus pose estimates were sufficiently accurate to generate meaningful latent space encodings and to produce interpretable density maps of distinct structures. It is clear, however, that this approach will fail if the degree of structural heterogeneity in the dataset results in inaccurate pose estimates. For example, a mixture of structurally unrelated complexes will align poorly to a consensus structure, and thus produce poor pose estimates. Notably, our framework is differentiable with respect to pose variables, which, in principle, should allow for on-the-fly pose-refinement or *de novo* pose estimation^25^, and future work will explore the efficacy of incorporating such features.

## Acknowledgments

The authors thank Ben Demeo, Ashwin Narayan, Adam Lerer, Roy Lederman, Sam Rodriques, Bob Sauer, Phil Sharp, and Kotaro Kelley for helpful discussions and feedback. This work was funded by the National Science Foundation Graduate Research Fellowship Program, NIH grant R01-GM081871 to BB, NIH grant R00-AG050749 to JD, and the MIT J-Clinic for Machine Learning and Health to JD and BB.

## Author Contributions

EZ, TB, BB, and JD conceived of the work. EZ, TB, and BB developed the representation learning method. EZ and JD tailored the method to cryo-EM data, and designed and analyzed the described experiments. EZ implemented the software and performed the experiments. EZ, JD, and BB wrote the manuscript.

## Methods

### The cryoDRGN method

#### Deep coordinate networks to represent 3D structure

The cryoDRGN method performs heterogeneous cryo-EM reconstruction by learning a neural network representation of 3D structure. In particular, we use a neural network to approximate the function *V*: ℝ^3+*n*^ → ℝ, which models structures as generated from an *n*-dimensional continuous latent space. We call this architecture^1-3^ a *deep coordinate network* as we explicitly model the volume as a function of Cartesian coordinates.

Without loss of generality, we model volumes on the domain [−0.5,0.5]^3^. Instead of directly supplying the 3D Cartesian coordinates, ***k***, to the deep coordinate network, coordinates are featurized with a fixed positional encoding function consisting of sinusoids whose wavelengths follow a geometric progression from 1 up to the Nyquist limit:

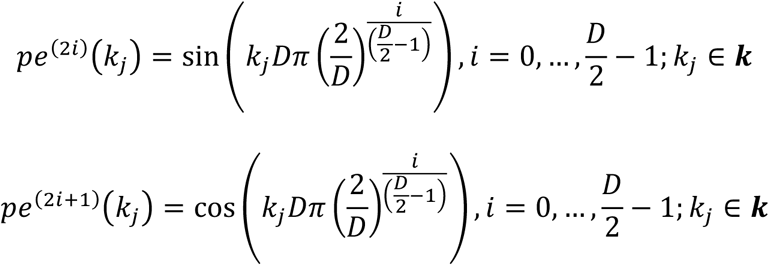

where *D* is set to the image size^1^ used in training. Empirically, we found that excluding the highest frequencies of the positional encoding led to better performance when training on noisy data, and we provide an option to modify the positional encoding function by increasing all wavelengths by a factor of 2*π*.

#### Training system

This parametric representation of 3D structure is learned via an image-encoder/volume-decoder architecture based on the variational autoencoder (VAE) ^4,5^. We follow the standard image formation model in single particle cryo-EM^3^ where observed images are generated from projections of a volume at a random unknown orientation, *R* ∈ *SO*(3). We use an additive Gaussian white noise model. Volume heterogeneity is generated from a continuous latent space, modeled by the latent variable ***z***, where the dimensionality of ***z*** is a hyperparameter of the model.

Given an image *X*, the variational encoder, *q*_*ξ*_(***z***|*X*), produces a mean and variance, *μ*_***z***|*X*_ and Σ_***z***|*X*_, statistics that parameterize a Gaussian distribution with diagonal covariance, as the variational approximation to the true posterior *p*(***z***|*X*). The prior on the latent variable is a standard normal distribution *𝒩*(0, **I**). The deep coordinate network architecture is used as the probabilistic decoder, *p*_*θ*_(*V*| ***k, z***), and models structures in frequency space. Given Cartesian coordinate ***k*** ∈ ℝ^3^ and latent variable ***z***, the probabilistic decoder predicts a Gaussian distribution over *V*(***k, z***). The encoder and decoder are parameterized with fully connected neural networks with parameters *ξ* and *θ*, respectively.

Since 2D projection images can be related to volumes as 2D central slices in Fourier space^6^, oriented 3D coordinates for a given image can be obtained by rotating a *D* × *D* lattice spanning [−0.5,0.5]^2^ originally on the x-y plane by *R*, the orientation of the volume during imaging. Then, given a sample out of *q*_*ξ*_(***z***|*X*) and the oriented coordinates, an image can be reconstructed pixel-by-pixel through the decoder. The reconstructed image is then translated by the image’s in-plane shift, and the CTF is applied before it is compared to the input image. The negative log likelihood of a given image under our model is computed as the mean square error between the reconstructed image and the input image. Following the standard VAE framework, the optimization objective is the variational lower bound of the model evidence:

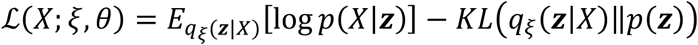

where the first term is the reconstruction error estimated with one Monte Carlo sample and the second term is a regularization term on the latent representation. By training on many 2D slices with sufficiently diverse orientations, the 3D volume can be learned through feedback from the 2D views. For further details, we refer the reader to a preliminary version of the method described in the proceedings of the International Conference for Learning Representations^3^. The results presented here employ the training regime described in Zhong *et al*. using previously determined poses from a consensus reconstruction^3^.

### Datasets

#### Simulated homogeneous dataset generation

The 50S subunit of the *E. coli* ribosome was extracted from PDB 4YBB in PyMOL^7^. A density map was generated from the atomic model using the molmap command in Chimera^8^ at a grid spacing of 1.5 Å/pix and a resolution of 4.5 Å. The resulting volume was padded to a box size of D=256, where D is the width in pixels along one dimension. Simulated micrographs were generated with custom Python scripts as follows: 50k projection images were generated by rotating the density map with a random rotation sampled uniformly from SO(3), projecting along the z-axis, and shifting the image with an in-plane translation sampled uniformly from [-20,20]^2^ pixels. Projection images were multiplied with the CTF in Fourier space, where the CTF was computed from defocus values randomly sampled from those given in EMPIAR-10028, no astigmatism, an accelerating voltage of 300 kV, a spherical aberration of 2mm, and an amplitude contrast ratio of 0.1. An envelope function with a B-factor of 100 Å^2^ was applied. Noise was added with a signal to noise ratio (SNR) of 0.1 where the noise-free signal images were defined as the entire DxD image. To generate the dataset with D=128, the D=256 noiseless projection images of the 50S were downsampled by Fourier clipping, followed by addition of CTF and noise as above.

#### Discrete3 heterogeneous dataset generation

To generate the “Discrete3” dataset, 10k, 15k, and 25k simulated micrographs of the 30S, 50S, and 70S ribosome, respectively, were combined. 15k micrographs from the homogeneous 50S dataset were used, and micrographs of the 30S and 70S ribosome were generated using the same procedure starting from the atomic model extracted from PDB 4YBB, and extracting either the 30S or 70S subunits. Images were downsampled to D=128, corresponding to a Nyquist limit of 6 Å.

#### Linear1D heterogeneous dataset generation

To generate the “Linear1D” dataset, 50 density maps were generated along a reaction coordinate defined by rotation of a dihedral angle in an atomic model of a hypothetical protein complex. Each model was generated at 0.03 radian increments of the bond rotation, leading to a total range of 1.5 radians. Density maps were generated in Chimera at a grid spacing of 6 Å/pix and resolution of 12 Å, and padded to a box size of D=128. 1000 projection images were generated with random orientations and in-plane translations from [-10,10]^2^ pixels for each map leading to a final particle stack of 50k images. CTF and noise at an SNR=0.1 were added using the same procedure described above.

#### Real cryo-EM datasets

Processed shiny particles and the star file containing CTF parameters were downloaded from the Electron Microscopy Public Image Archive (EMPIAR) ^9^ for datasets EMPIAR-10028, EMPIAR-10076, and EMPIAR-10180. Particle images were resized to either D=96, 128, or 256 by clipping in Fourier space with a custom Python script. These various images sizes resulted in the following Nyquist limits:

| Dataset name | EMPIAR ID | Image size, D (pixels) | Nyquist limit (Å) | Figure |
| --- | --- | --- | --- | --- |
| 80S ribosome | 10028 | 96 | 10.1 | N/A |
| 80S ribosome | 10028 | 256 | 3.8 | 4 |
| Assembling LSU ribosome | 10076 | 128 | 6.6 | 5 |
| Assembling LSU ribosome | 10076 | 256 | 3.3 | 5 |
| Pre-catalytic spliceosome | 10180 | 128 | 8.5 | 6 |
| Pre-catalytic spliceosome | 10180 | 256 | 4.3 | 6 |

### Traditional homogeneous reconstruction

3D reconstruction of the 80S ribosome (EMPIAR-10028) was performed in cryoSPARC v2.4^10^ using the ab-initio reconstruction job followed by the homogeneous refinement job with default parameters. The final reconstruction reported a GSFSC_0.143_^11^ resolution of 3.1 Å with a tight mask and 4.1 Å unmasked. The density map was sharpened using the published B-factor of -80.1 Å^2^ for visualization.

Homogeneous 3D reconstruction of the L17-depleted ribosome assembly intermediates (EMPIAR 10076) was performed as above, leading to a final structure with a GSFSC_0.143_^11^ resolution of 3.2 Å with a tight mask and 4.8 Å unmasked.

### Deep coordinate network training on homogeneous structures

For each dataset and for each architecture, a separate deep coordinate network with no latent variable was trained for 25 epochs, where an epoch is defined as one pass through the dataset. The tested architectures were fully connected networks with ReLU activations, where the network size was either 3 layers of dimension 128 (128 nodes/layer x 3 layers), 3 layers of dimension 256 (256×3), 3 layers of dimension 1024 (1024×3), or 10 layers of dimension 1024 (1024×10). Image poses were set to either the ground truth poses for the simulated datasets, or poses obtained from a traditional homogeneous reconstruction in cryoSPARC. Networks were trained on minibatches of 8 images using the Adam^12^ optimizer with a learning rate of 0.0001. Once training completed, the deep coordinate network was evaluated on the 3D coordinates of a *D* × *D* × *D* voxel array spanning [-0.5,0.5]^3^, where D is the image size in pixels along one dimension. The density map was sharpened using the published B-factor of -80.1 Å^2^ for visualization^13^.

### Map-to-map FSC

Fourier shell correlation curves were computed between the ground truth density maps and the neural network reconstructed density maps using a custom Python script. For the homogeneous reconstruction of EMPIAR 10028, the map-to-map FSC was computed between the neural network structure and the traditional homogeneous reconstruction in cryoSPARC after applying a real space mask and with phase randomization at frequencies above 3.1 Å, the GSFSC_0.143_ of the cryoSPARC reconstruction. The real space mask was defined by first thresholding the volume at half of the 99.99th percentile density value. The mask was then dilated by 15 Å from the original boundary, and a soft cosine edge was used to taper the mask to 0 at 25 Å from the original boundary.

### CryoDRGN heterogeneous reconstruction

CryoDRGN encoder-decoder networks were trained from their randomly initialized values for each single particle cryo-EM dataset. Unless otherwise specified, all networks were trained on minibatches of 8 images using the Adam optimizer with a learning rate of 0.0001. After training, the dataset was evaluated through the encoder, and the *maximum a posteriori* value of *q*(***z***|*X*) was defined as the latent encoding for each image. Visualization of the latent encodings with PCA and UMAP and analysis with k-means clustering was performed with scikit-learn^14^. Density maps were generated by evaluating the decoder on a desired value of the latent variable *z* and the 3D coordinates of a *D* × *D* × *D* voxel array spanning [-0.5,0.5]^3^.

#### Heterogeneous reconstruction of simulated datasets

For each simulated heterogeneous dataset, a 1D latent variable model was trained for 100 epochs. The encoder architecture was 256×3 (nodes/layer x layers) and the decoder architecture was 512×5. The image poses used for training were the ground truth image poses. Structures shown in Figure 3b were generated at the 5th, 23rd, 41st, 59th, 77th, and 95^th^ percentile values of the latent encodings, and sharpened by a B-factor of -100 Å^2^. Structures shown in Figure 3e were generated at the *k*-means cluster centers after performing *k*-means clustering with *k*=3 on the latent encodings, and sharpened by a B-factor of -100 Å^2^.

#### Heterogeneous reconstruction of the 80S ribosome (EMPIAR-10028)

##### Pilot experiments

A 10D latent variable model was trained on downsampled images (D=96, 4.91Å/pix) from EMPIAR-10028 for 50 epochs. The encoder and decoder architectures were 128×10, and the mini-batch size was 5. Image poses were obtained from a traditional homogeneous reconstruction in cryoSPARC.

##### Particle filtering

After training, *k*-means clustering with *k*=20 was performed on the predicted latent encodings for the dataset. One cluster contained 860 particles that were outliers when viewing the projected encodings along the first and second principle component. This observation was reproducible, and the particles belonging to the outlier cluster from either of two replicates (960 particles in total) were removed from the dataset.

##### High resolution training

After particle filtering, a 10D latent variable model was trained on the remaining 104,280 images (D=256, 1.84 Å/pix) for 150 epochs. The encoder and decoder architectures were 1024×3.

##### Analysis

After training, *k*-means clustering with *k*=20 was performed on the predicted latent encodings for the dataset, and volumes were generated at the cluster centers using the decoder network. Representative structures were manually selected for visualization in Figure 4.

#### Heterogeneous reconstruction of the L17-depleted ribosome assembly intermediates (EMPIAR-10076)

##### Pilot experiments

A 10D latent variable model was trained on downsampled images (D=128, 3.3 Å/pix) from EMPIAR 10076 for 50 epochs. The encoder and decoder architectures were 256×3. Image poses were obtained from a traditional homogeneous reconstruction in cryoSPARC.

##### Particle filtering

Particles with ***z*** ≤ −1 were removed from subsequent analysis.

##### High resolution training

A 10D latent variable model was trained on the remaining 101,604 images (D=256, 1.7 Å/pix) for 50 epochs. The encoder and decoder architectures were 1024×3.

##### Analysis

After training, the dataset’s latent encodings were viewed in 2D with UMAP^15^. Density maps corresponding to the major and minor assembly states were generated at the mean latent encoding for each class, *i*.*e*. 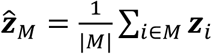, where M is the set of particles assigned to a given class in the published 3D classification. Instead of evaluating the volume decoder at 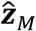, we find the latent encoding of the dataset closest in Euclidean distance to 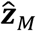 as the “on data” representative encoding.

##### New assembly state

Particles corresponding to the new assembly state (C4) were manually selected from the UMAP embeddings with an interactive lasso tool in a custom visualization script. The mean latent encoding of the resulting 1,211 selected particles was used to generate the structure representative for this new assembly state.

##### Voxel-based back-projection

The particles associated with class C4 and their corresponding poses were used to reconstruct a structure via traditional voxel-based back-projection using a custom Python script. In this simplified implementation, images were first phase flipped to correct for the CTF. Then each image was centered by its in-plane translation and aligned in 3D space based on its 3D rotation. The density for each voxel was computed using a linear interpolation kernel. The structure was then low-pass filtered to 8 Å for visual clarity.

#### Heterogeneous reconstruction of the pre-catalytic spliceosome (EMPIAR-10180)

##### Pilot experiments

A 10D latent variable model was trained on downsampled images (D=128, 4.25 Å/pix) from EMPIAR 10180 for 50 epochs. The encoder and decoder architectures were 256×3. Poses were obtained from the consensus reconstruction values given in the consensus_data.star deposited to EMPIAR 10180.

##### Particle filtering

The UMAP embeddings showed multiple clusters where the largest cluster corresponded to fully formed pre-catalytic spliceosomes. Particles corresponding to other clusters were removed from subsequent analyses by first performing *k*-means clustering with *k*=20 on the latent encodings, and removing *k*-means clusters whose structure did not resemble the fully formed pre-catalytic spliceosome (11 out of 20 *k*-means clusters in one replicate, and 10 out of 20 in a second replicate).

##### High resolution training

A 10D latent variable model was trained on the remaining 150,098 images (D=256, 2.1 Å/pix) for 50 epochs. The encoder and decoder architectures were 1024×3.

##### Analysis

After training, the dataset’s latent encoding was viewed in 2D with UMAP (Fig. 6a) and PCA (Fig. 6c). Density maps in Figure 6d were generated at the latent encoding values that traverse PC1 at five equally spaced points between the 5^th^ and 95^th^ percentile of PC1 values. Density maps in Extended Fig. 7 were generated at the latent encoding values that traverse PC2 at five equally spaced points between the 5^th^ and 95^th^ percentile of PC2 values.

### Latent space graph traversal for generating trajectories

Trajectories were generated by first creating a nearest-neighbors graph from the latent encodings of the images, where a neighbor was defined if the Euclidean distance was below a threshold computed from the statistics of all pairwise distances. We choose a value such that the average number of neighbors across all nodes is 5. Edges were then pruned such that a given node does not have more than 10 neighbors. Then, Djikstra’s algorithm was used to find the shortest path along the graph connecting a series of anchor points, and density maps were generated at the ***z*** value of the visited nodes. Anchor points were set to be the “on-data” cluster centers after performing *k*-means clustering of the latent encodings with *k*=20. Instead of using the mean value of each *k*-means cluster, we define the latent encoding closest in Euclidean distance to the k-means cluster center as the “on-data” cluster center.

To generate Supplemental Movie 1 of the 80S ribosome, 113 density maps were generated by following the protocol above, and we visualized a representative sequence of 60 density maps that contained the 40S rotated state. To generate Supplemental Movies 2 and 3 of the assembling bacterial LSU, anchor points were manually chosen from an interactive tool provided in cryoDRGN to create a path along the C-class assembly pathway and the D-class assembly pathway. To generate Supplemental Movie 4 of the pre-catalytic spliceosome, 132 density maps were generated following the above protocol.

### 2D class averages

2D classification was performed in cryoSPARC^10^ using all default options except for the number of 2D classes, which was set to 20.

### Data availability

Trained cryoDRGN models for all experiments, simulated datasets, and indices of filtered particles of EMPIAR-10028, EMPIAR-10076, and EMPIAR-10180 are available upon request.

### Software availability

All software and analysis scripts are implemented in custom Python code using PyTorch^16^ and are available at cryodrgn.csail.mit.edu.

---

**Supplementary Figure 1.**
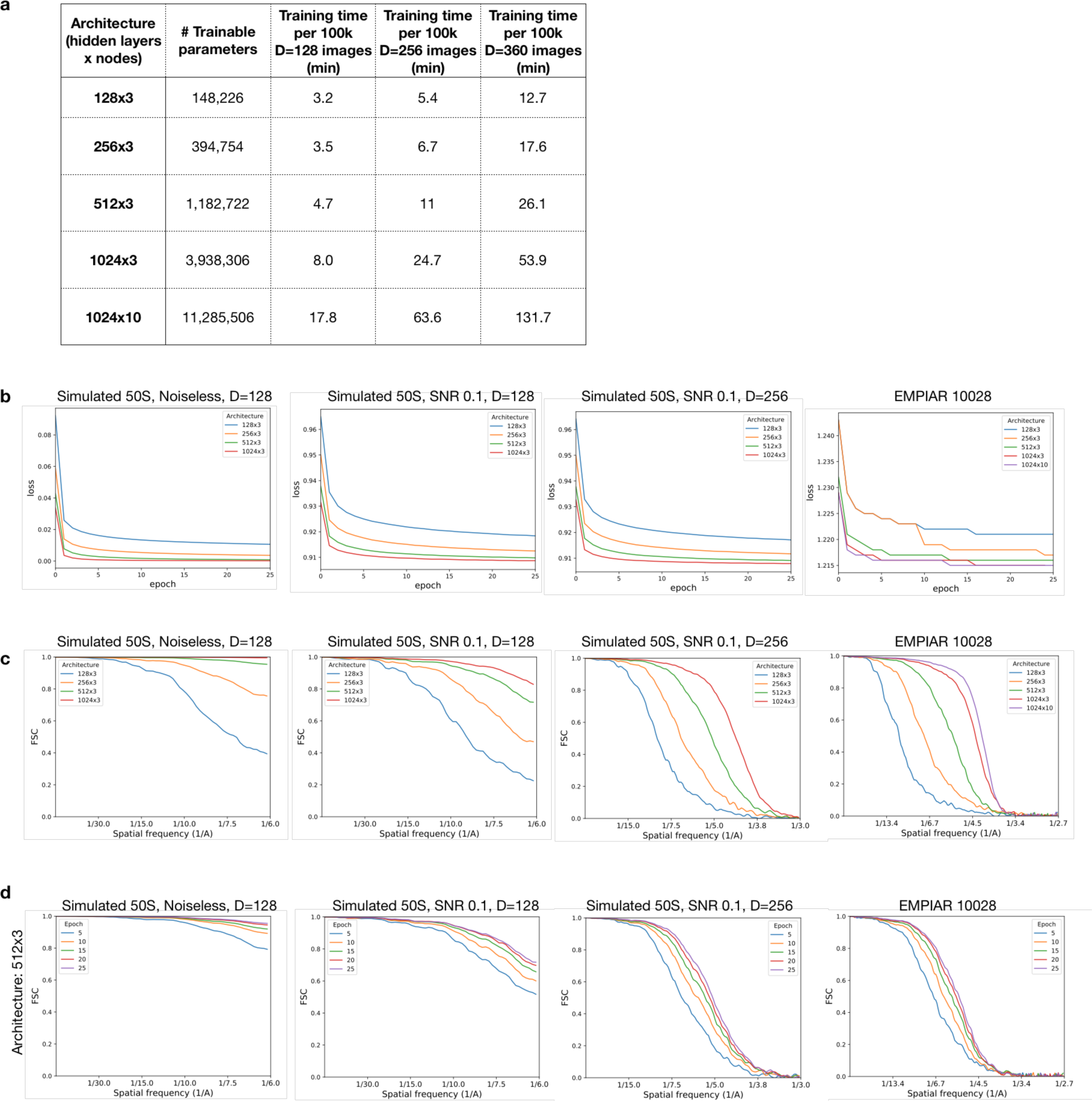
Neural network training statistics for homogeneous reconstruction. **a)** Training time in minutes per 100k images for different architectures and image sizes on a single Nvidia V100 GPU. **b)** Loss curve for training deep coordinate networks of varying architectures on four different datasets: 50k simulated noiseless projection images of the 50S ribosome (D=128, Nyquist limit of 6 Å), 50k simulated micrographs of the 50S ribosome (D=128), 50k simulated micrographs of the 50S ribosome (D=256, Nyquist limit of 3 Å), and 104,249 micrographs of the 80S ribosome from EMPIAR 10028 (D=360, Nyquist limit of 2.68 Å). **c)** FSC curve between the ground truth density map and the learned density map after 25 epochs of training deep coordinate networks of varying architectures. **d)** FSC curve between the ground truth density map and the learned density map at different epochs of training a deep coordinate network with 3 hidden layers of dimension 512. We use the traditionally reconstructed map as the ground truth structure for EMPIAR 10028.

**Supplementary Figure 2.**
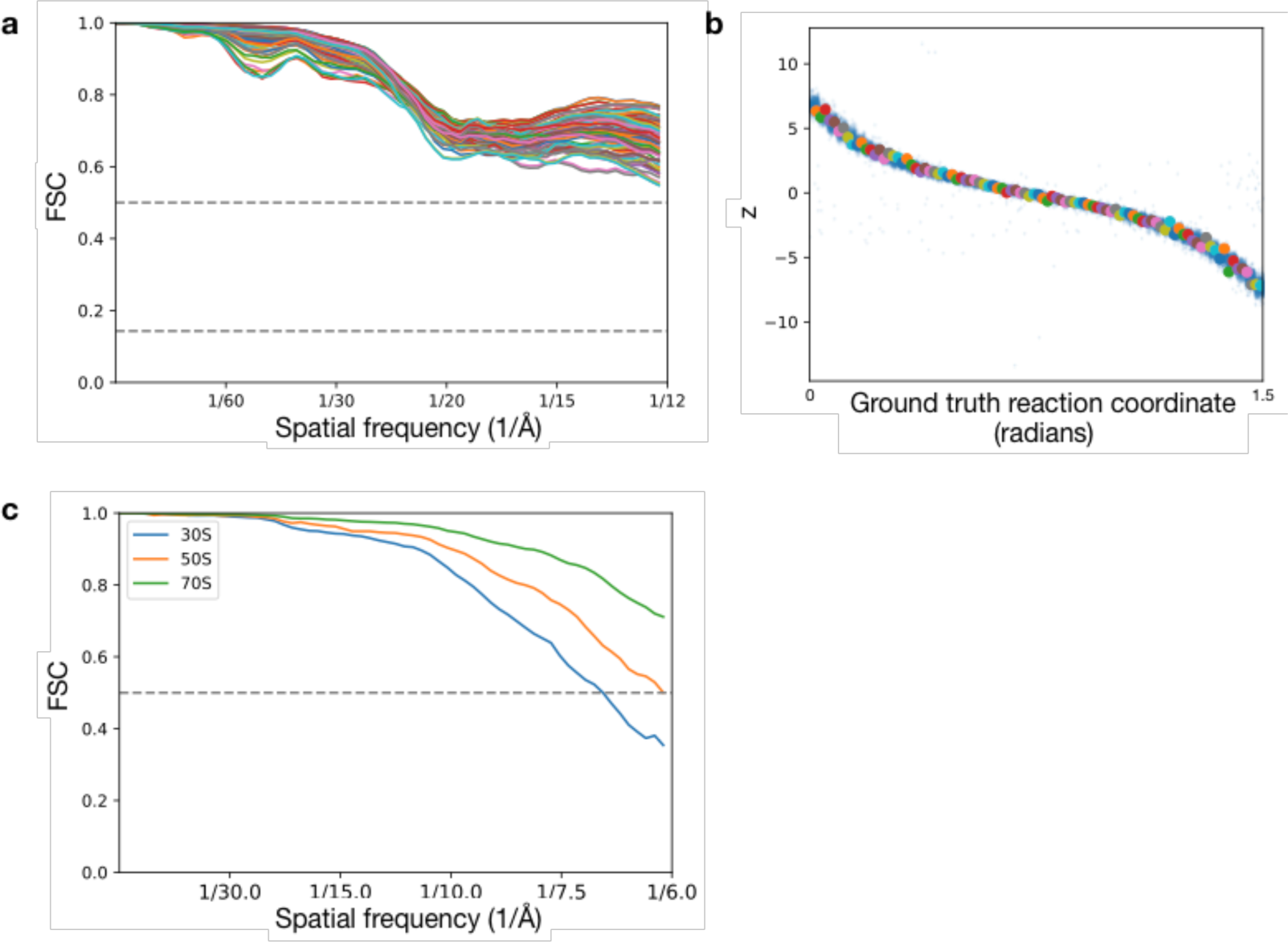
FSC curves between ground truth maps and density maps from cryoDRGN trained on heterogeneous simulated datasets. **a)** 100 FSC curves between generated and ground truth density maps. The density maps are generated at the value of the latent variable predicted for a given image, and compared against the ground truth density map that generated the image. Images are uniformly sampled along the reaction coordinate. **b)** The predicted latent encoding for the 100 images along the ground truth reaction coordinate for the density maps in (a). **c)** FSC curve between the generated density maps shown in Figure 3f and the ground truth 30S, 50S, and 70S density map.

**Supplementary Figure 3.**
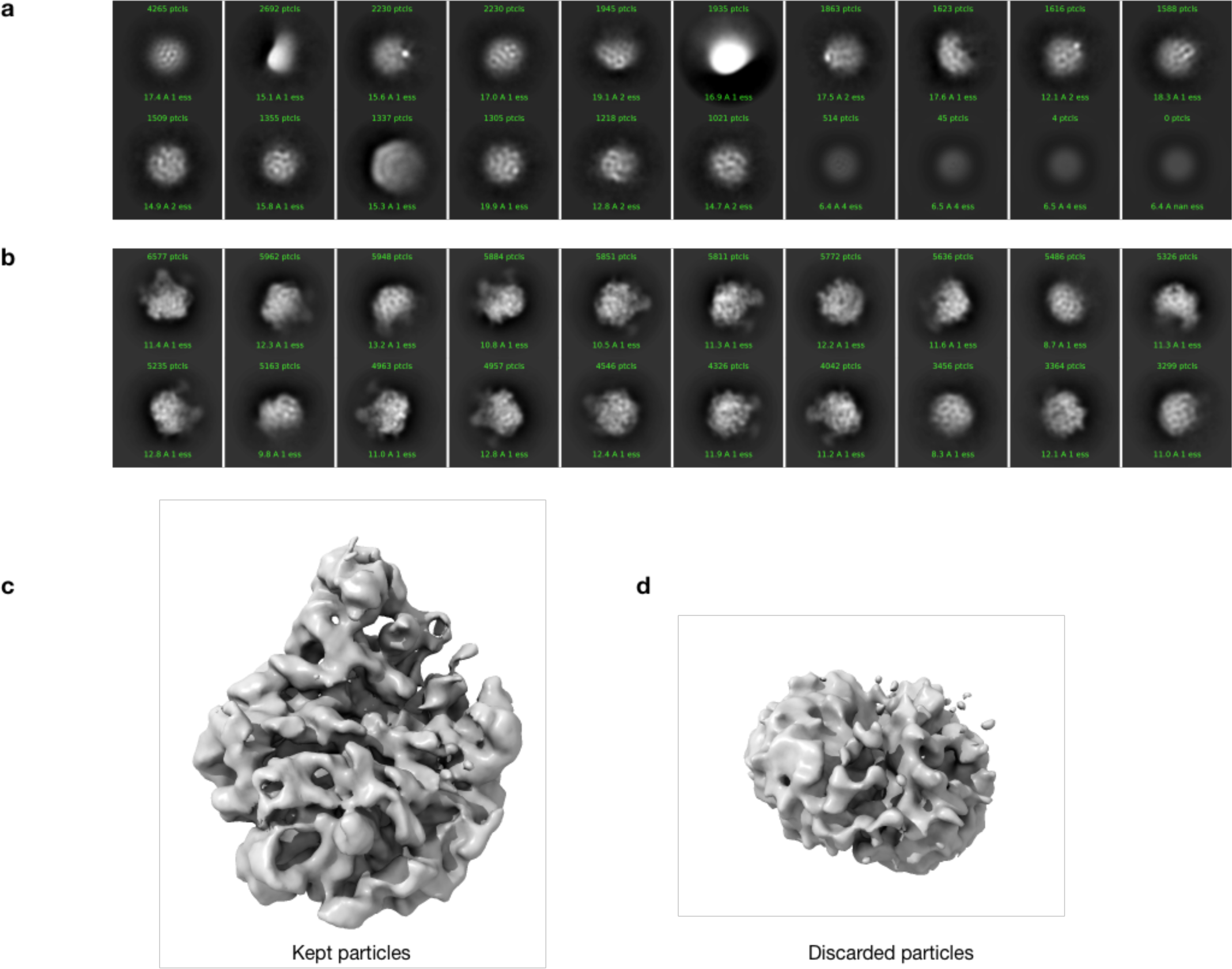
Filtering of particles from the assembling ribosome dataset. **a)** 2D class averages of discarded particles with *z* ≤ − 1 from Figure 5a. **b)** 2D class averages of kept particles with *z* > 1 from Figure 5a. **c)** CryoSPARC *ab-initio* reconstruction of kept particles (*z* > −1) and **d)** of discarded particles (*z* ≤ − 1) from Figure 5a.

**Supplementary Figure 4.**
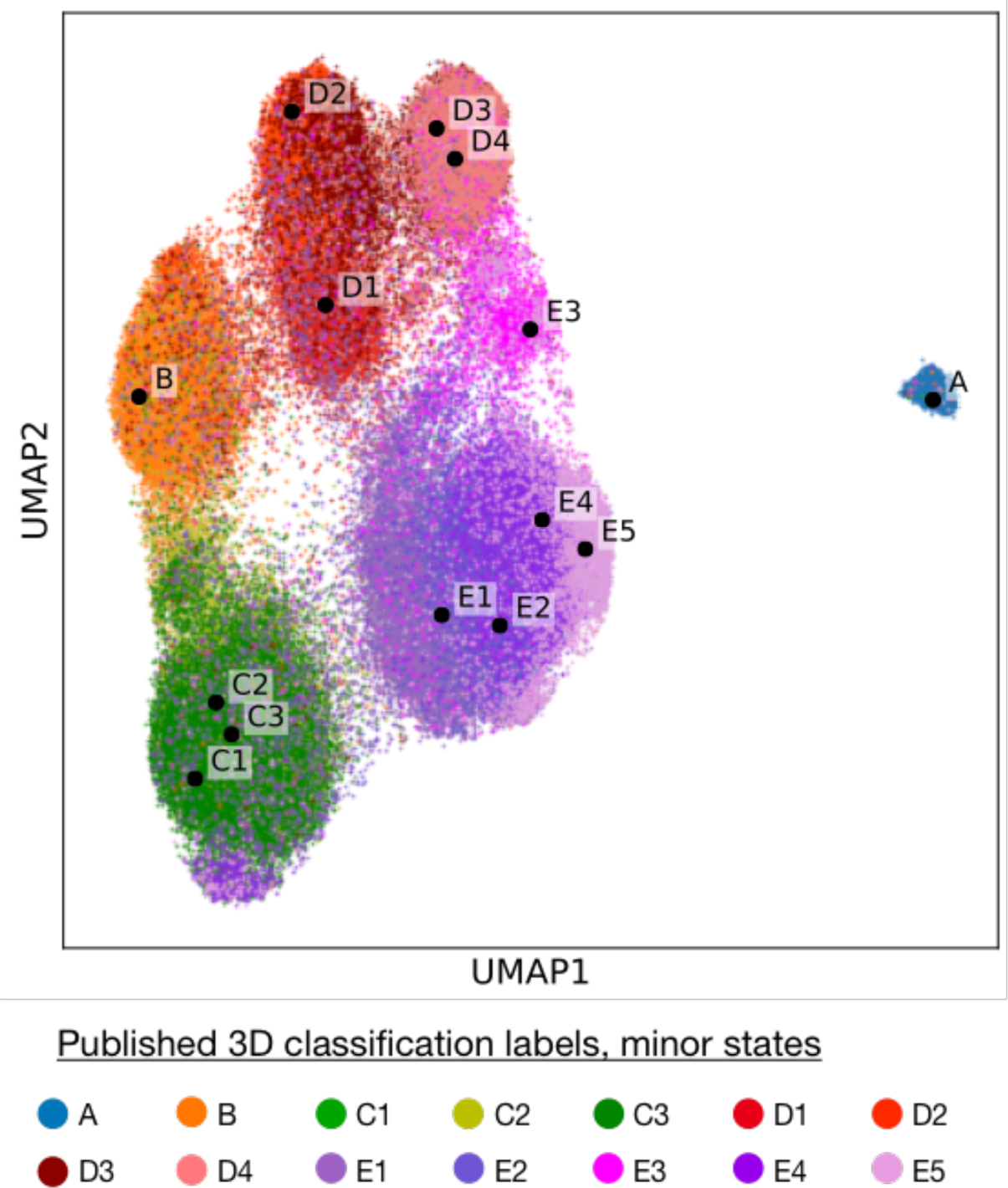
CryoDRGN latent encodings trained on the assembling ribosome. UMAP embedding of the latent space encodings of particle images after training a cryoDRGN 10D latent variable model on EMPIAR 10076. Points are colored by the 3D classification labels corresponding to the minor states of LSU assembly from *Davis et al*.

**Supplementary Figure 5.**
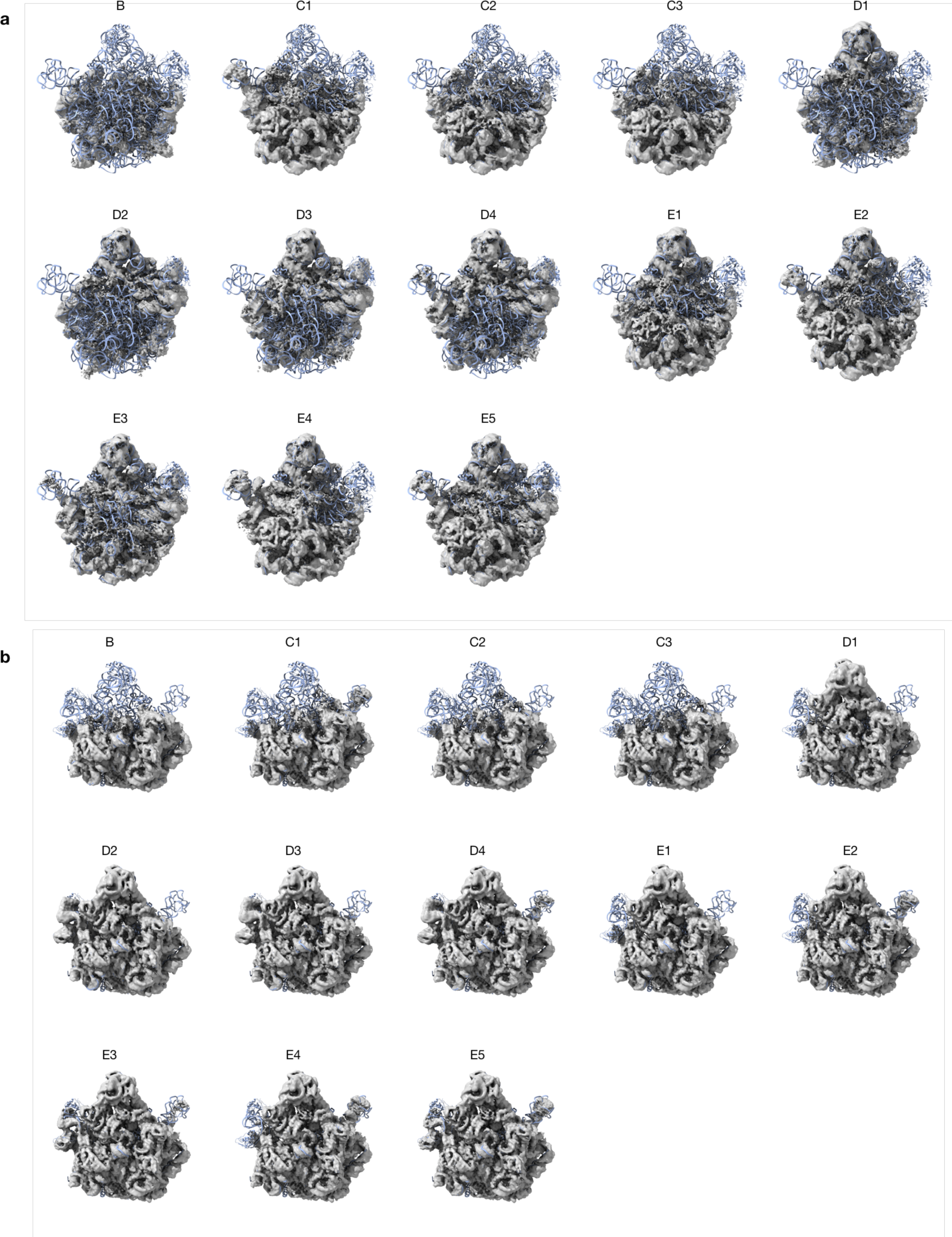
Minor LSU assembly states reconstructed from cryoDRGN trained on the assembling ribosome dataset. **a)** Front view and **b)** back view of minor state density maps after training a cryoDRGN 10D latent variable model on particle images from EMPIAR 10076 with the 50S crystal structure docked (PDB 4YBB). Each cryoDRGN structure is generated at the latent variable values shown in Supplementary Figure 4, which are computed from the mean latent encoding of particles with the corresponding class assignment from *Davis et al*. Views match perspectives from Figure 5d.

**Supplementary Figure 6.**
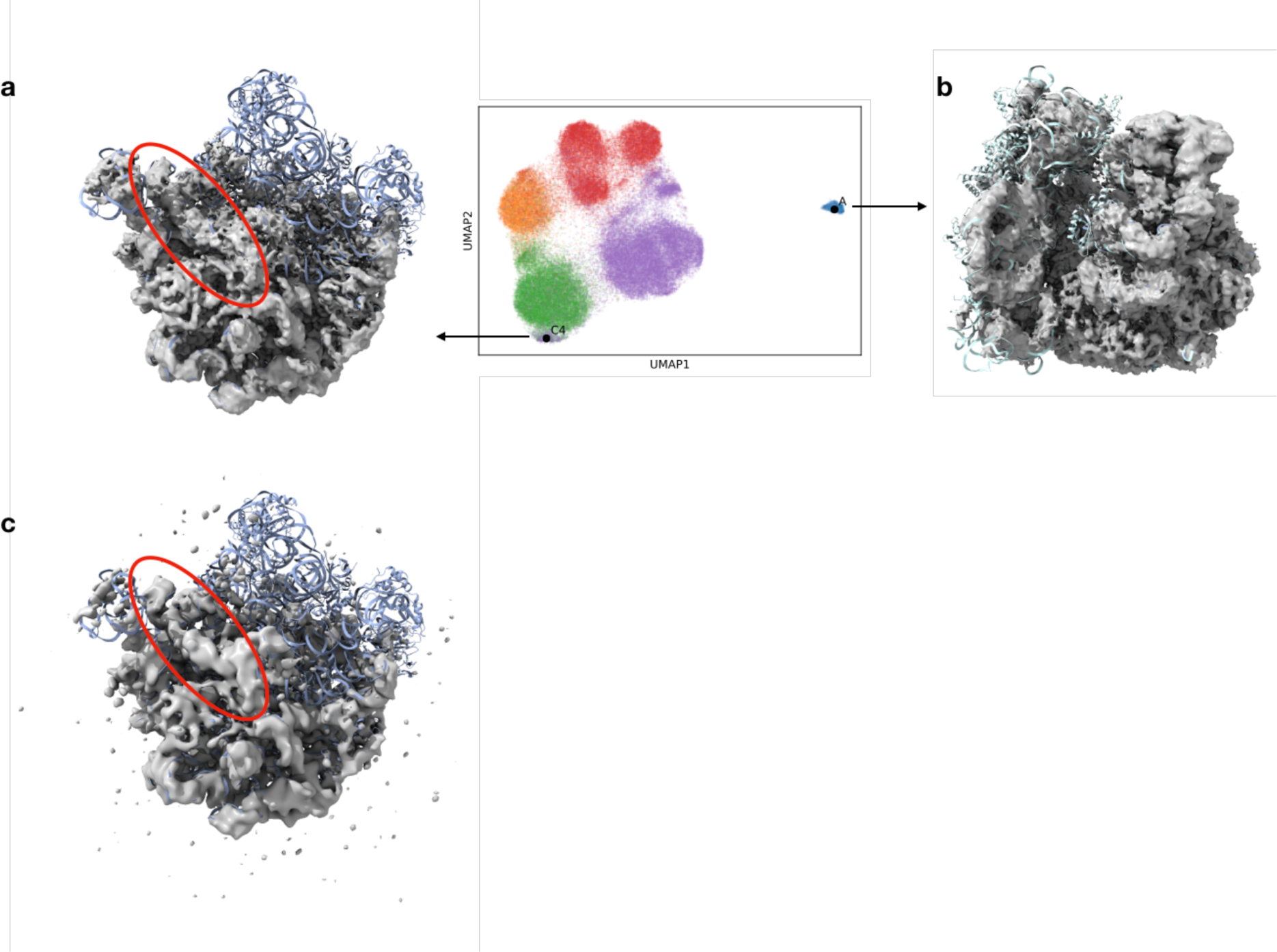
Additional structures reconstructed from cryoDRGN trained on the assembling ribosome dataset. **a)** Density map of a new assembly state, class C4, produced by cryoDRGN. Helix 68 (red oval) was exclusively associated with mature classes E4 and E5 in Davis *et al*. The structure is generated from point C4 in latent space, which belongs to a small cluster proximal to class C that was classified into class E4 and E5 by Davis *et al*. **b)** The density map of the 70S ribosome reconstructed by cryoDRGN from point A in latent space. **c)** Voxel-array backprojection of the 1,211 particles contained in the latent space cluster corresponding to the new assembly state with atomic model docked and helix 68 highlighted (red oval).

**Supplementary Figure 7.**
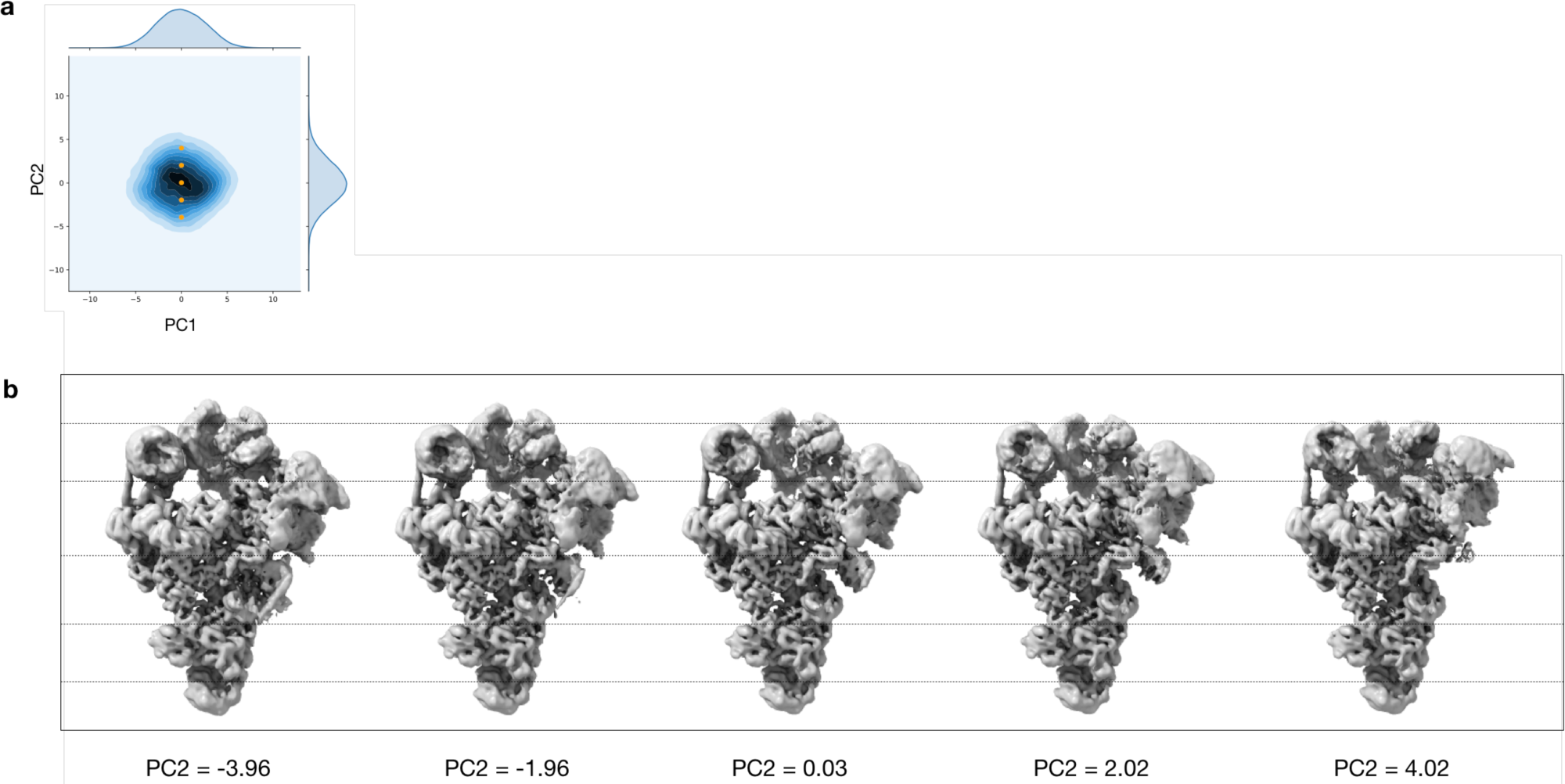
Additional structures of the pre-catalytic spliceosome reconstructed by cryoDRGN. **a)** PCA projections of the 10D latent encodings from cryoDRGN with 5 points along PC2 shown in orange. b) Density maps produced by cryoDRGN at the 5 highlighted points from (a).

**Supplemental Movie 1**. Trajectory of the pre-catalytic spliceosome. Traversal in latent space (left) and corresponding structures generated from cryoDRGN (right).

**Supplemental Movie 2**. Trajectory of the assembling ribosome along the D-class assembly pathway described in Davis *et al*. Traversal in latent space (left) and corresponding structures generated from cryoDRGN (right).

**Supplemental Movie 3**. Trajectory of the assembling ribosome along the C-class assembly pathway described in Davis *et al*. Traversal in latent space (left) and corresponding structures generated from cryoDRGN (right).

**Supplemental Movie 4**. Trajectory of the 80S ribosome. Traversal in latent space (left) and corresponding structures generated from cryoDRGN (right).

## Footnotes

1 Number of pixels along one dimension of the image, i.e. a *D* × *D* image

## References

1. Nogales, E. The development of cryo-EM into a mainstream structural biology technique. Nat. Methods 13, 24–27 (2016).

2. Cheng, Y. Single-particle cryo-EM—How did it get here and where will it go. Science 361, 876–880 (2018).

3. Bammes, B. E., Rochat, R. H., Jakana, J., Chen, D.-H. & Chiu, W. Direct electron detection yields cryo-EM reconstructions at resolutions beyond 3/4 Nyquist frequency. J. Struct. Biol. 177, 589–601 (2012).

4. Suloway, C. et al. Automated molecular microscopy: The new Leginon system. J. Struct. Biol. 151, 41–60 (2005).

5. Li, X. et al. Electron counting and beam-induced motion correction enable near-atomic-resolution single-particle cryo-EM. Nat. Methods 10, 584–590 (2013).

6. Zhang, K. Gctf: Real-time CTF determination and correction. J. Struct. Biol. 193, 1–12 (2016).

7. Brubaker, M. A., Punjani, A. & Fleet, D. J. Building Proteins in a Day: Efficient 3D Molecular Reconstruction. 3099–3108 (2015).

8. Scheres, S. H. W. A Bayesian view on cryo-EM structure determination. J. Mol. Biol. 415, 406–418 (2012).

9. Bepler, T. et al. Positive-unlabeled convolutional neural networks for particle picking in cryo-electron micrographs. Nat Methods 16, 1153–1160 (2019).

10. Ahmed, T., Yin, Z. & Bhushan, S. Cryo-EM structure of the large subunit of the spinach chloroplast ribosome. Sci Rep 6, 1–13 (2016).

11. Wrapp, D. et al. Cryo-EM structure of the 2019-nCoV spike in the prefusion conformation. Science 367, 1260–1263 (2020).

12. Sigworth, F. J. Principles of cryo-EM single-particle image processing. Microscopy (Oxf) 65, 57–67 (2016).

13. Scheres, S. H. W. et al. Maximum-likelihood Multi-reference Refinement for Electron Microscopy Images. J. Mol. Biol. 348, 139–149 (2005).

14. Lyumkis, D., Brilot, A. F., Theobald, D. L. & Grigorieff, N. Likelihood-based classification of cryo-EM images using FREALIGN. J. Struct. Biol. 183, 377–388 (2013).

15. Scheres, S. H. W. et al. Disentangling conformational states of macromolecules in 3D-EM through likelihood optimization. Nat. Methods 4, 27–29 (2007).

16. Haselbach, D. et al. Structure and Conformational Dynamics of the Human Spliceosomal Bact Complex. Cell 172, 454–464.e11 (2018).

17. Nakane, T., Kimanius, D., Lindahl, E. & Scheres, S. H. Characterisation of molecular motions in cryo-EM single-particle data by multi-body refinement in RELION. Elife 7, e36861 (2018).

18. Frank, J. & Ourmazd, A. Continuous changes in structure mapped by manifold embedding of single-particle data in cryo-EM. Methods 100, 61–67 (2016).

19. Moscovich, A., Halevi, A., Andén, J. & Singer, A. Cryo-EM reconstruction of continuous heterogeneity by Laplacian spectral volumes. arXiv.org eess.IV, (2019).

20. Lederman, R. R. & Singer, A. Continuously heterogeneous hyper-objects in cryo-EM and 3-D movies of many temporal dimensions. arXiv.org cs.CV, (2017).

21. Iudin, A., Korir, P. K., Salavert-Torres, J., Kleywegt, G. J. & Patwardhan, A. EMPIAR: a public archive for raw electron microscopy image data. Nat. Methods 13, 387–388 (2016).

22. GPLv3 GNU General Public License. Free Software Foundation (2007).

23. Bricman, P. A. & Ionescu, R. T. CocoNet: A deep neural network for mapping pixel coordinates to color values. arXiv.org cs.CV, (2018).

24. Bepler, T., Zhong, E., Kelley, K., Brignole, E. & Berger, B. Explicitly disentangling image content from translation and rotation with spatial-VAE. Advances in Neural Information Processing Systems 32 15435–15445 (2019).

25. Zhong, E. D., Bepler, T., Davis, J. H. & Berger, B. Reconstructing continuous distributions of 3D protein structure from cryo-EM images. In International Conference on Learning Representations (2020).

26. Kingma, D. P. & Welling, M. Auto-Encoding Variational Bayes. arXiv.org stat.ML, (2013).

27. Rezende, D. J., Mohamed, S. & Wierstra, D. Stochastic Backpropagation and Approximate Inference in Deep Generative Models. arxiv.org (2014).

28. Bracewell, R. N. Strip Integration in Radio Astronomy. Aust. J. Phys. 9, 198–217 (1956).

29. Wong, W. et al. Cryo-EM structure of the Plasmodium falciparum 80S ribosome bound to the anti-protozoan drug emetine. Elife 3, e01963 (2014).

30. Punjani, A., Rubinstein, J. L., Fleet, D. J. & Brubaker, M. A. cryoSPARC: algorithms for rapid unsupervised cryo-EM structure determination. Nat. Methods 14, 290–296 (2017).

31. Sun, M. et al. Dynamical features of the Plasmodium falciparum ribosome during translation. Nucleic Acids Research 2;43(21):10515–24 (2015).

32. Davis, J. H. et al. Modular Assembly of the Bacterial Large Ribosomal Subunit. Cell 167, 1610–1622.e15 (2016).

33. McInnes, L., Healy, J. & Melville, J. UMAP: Uniform Manifold Approximation and Projection for Dimension Reduction. arxiv.org (2018).

34. Plaschka, C., Lin, P.-C. & Nagai, K. Structure of a pre-catalytic spliceosome. Nature 546, 617–621 (2017).

35. Hornik, K. Approximation capabilities of multilayer feedforward networks. Neural Networks 4, 251–257 (1991).

36. Cormen, Thomas H.; Leiserson, Charles E.; Rivest, Ronald L.; Stein, Clifford (2001). “Section 24.3: Dijkstra’s algorithm”. Introduction to Algorithms (Second ed.). MIT Press and McGraw–Hill. pp. 595–601. ISBN 0-262-03293-7.

37. Zhang, C., Bengio, S., Hardt, M., Recht, B. & Vinyals, O. Understanding deep learning requires rethinking generalization. arxiv.org (2016).

38. Buhai, R.-D., Risteski, A., Halpern, Y. & Sontag, D. Benefits of Overparameterization in Single-Layer Latent Variable Generative Models. (2019).

39. Zivanov, J. et al. RELION-3: new tools for automated high-resolution cryo-EM structure determination. bioRxiv 421123 (2018). doi: 10.1101/421123

40. Punjani, A., Zhang, H. & Fleet, D. J. Non-uniform refinement: Adaptive regularization improves single particle cryo-EM reconstruction. bioRxiv 179, 2019.12.15.877092 (2019).

## References

1. Bricman, P. A. & Ionescu, R. T. CocoNet: A deep neural network for mapping pixel coordinates to color values. arXiv.org cs.CV, (2018).

2. Bepler, T., Zhong, E., Kelley, K., Brignole, E. & Berger, B. Explicitly disentangling image content from translation and rotation with spatial-VAE. 15435–15445 (2019).

3. Zhong, E. D., Bepler, T., Davis, J. H. & Berger, B. Reconstructing continuous distributions of 3D protein structure from cryo-EM images. arXiv.org q-bio.QM, (2019).

4. Kingma, D. P. & Welling, M. Auto-Encoding Variational Bayes. arXiv.org stat.ML, (2013).

5. Rezende, D. J., Mohamed, S. & Wierstra, D. Stochastic Backpropagation and Approximate Inference in Deep Generative Models. arxiv.org (2014).

6. Bracewell, R. N. Strip Integration in Radio Astronomy. Aust. J. Phys. 9, 198–217 (1956).

7. The PyMOL Molecular Graphics System, Version 2.3 Schrödinger, LLC.

8. Pettersen, E. F. et al. UCSF Chimera—A visualization system for exploratory research and analysis. Journal of Computational Chemistry 25, 1605–1612 (2004).

9. Iudin, A., Korir, P. K., Salavert-Torres, J., Kleywegt, G. J. & Patwardhan, A. EMPIAR: a public archive for raw electron microscopy image data. Nat. Methods 13, 387–388 (2016).

10. Punjani, A., Rubinstein, J. L., Fleet, D. J. & Brubaker, M. A. cryoSPARC: algorithms for rapid unsupervised cryo-EM structure determination. Nat. Methods 14, 290–296 (2017).

11. Rosenthal, P. B. & Henderson, R. Optimal determination of particle orientation, absolute hand, and contrast loss in single-particle electron cryomicroscopy. J. Mol. Biol. 333, 721–745 (2003).

12. Kingma, D. P. & Ba, J. Adam: A Method for Stochastic Optimization. arxiv.org (2014).

13. Wong, W. et al. Cryo-EM structure of the Plasmodium falciparum 80S ribosome bound to the anti-protozoan drug emetine. Elife 3, e01963 (2014).

14. Pedregosa, F. et al. Scikit-learn: Machine Learning in Python. Journal of Machine Learning Research 12, 2825–2830 (2011).

15. McInnes, L., Healy, J. & Melville, J. UMAP: Uniform Manifold Approximation and Projection for Dimension Reduction. arxiv.org (2018).

16. Paszke, A. et al. PyTorch: An Imperative Style, High-Performance Deep Learning Library. 8026–8037 (2019).

